# An integrated genome-wide multi-omics analysis of gene expression dynamics in the preimplantation mouse embryo

**DOI:** 10.1101/495788

**Authors:** Steffen Israel, Mathias Ernst, Olympia E. Psathaki, Hannes C. A. Drexler, Ellen Casser, Yutaka Suzuki, Wojciech Makalowski, Michele Boiani, Georg Fuellen, Leila Taher

## Abstract

Early mouse embryos have an atypical translational machinery comprised of cytoplasmic lattices, poorly competent for translation. Thus, the impact of transcriptomic changes on the operational levels of proteins has likely been overestimated in the past. To find out, we used liquid chromatography–tandem mass spectrometry to detect and quantify 6,550 proteins in the oocyte and in six developmental stages (from zygote to blastocyst) collected in triplicates, and we also performed mRNA sequencing.

In contrast to the known split between the 2-cell and 4-cell stages at the transcript level, on the protein level the oocyte-to-embryo transition appeared to last until the morula stage. In general, protein abundance profiles were weakly correlated with those of their cognate mRNAs and we found little or no concordance between changes in protein and transcript expression relative to the oocyte at early stages. However, concordance increased towards morula and blastocyst, hinting at a more direct coupling of proteins with transcripts at these stages, in agreement with the increase in free ribosome abundance. Independent validation by immunofluorescence and qPCR confirmed the existence of genes featuring strongly positively and negatively correlated protein and transcript. Moreover, consistent coverage of most known protein complexes indicates that our dataset represents a large fraction of the expressed proteome. Finally, we identified 20 markers, including members of the endoplasmic reticulum pathway, for discriminating between early and late stages.

This resource contributes towards closing the gap between the ‘predicted’ phenotype, based on mRNA, and the ‘actual’ phenotype, based on protein, of the mouse embryo.

## Introduction

It has been about *100 years* since the *mouse* became a premier *model* organism. This status has been reinforced by the arrival of high-throughput RNA sequencing technologies, making it possible to investigate the regulatory circuits underlying development in detail. However, it is uncertain how closely RNA changes correlate with the operational level of the proteins. In fact, work in plants, yeast, lower vertebrates (Smits et al. 2014) and mammalian cell lines (Schwanhausser et al. 2011) has revealed a modest correlation. Mouse oocytes and early embryos feature an atypical translational machinery regarded to be poorly competent for mRNA translation (‘cytoplasmic lattices’ in place of free ribosomes, (Yurttas et al. 2008)). Thus, the impact of transcriptional changes on the embryo proteome is expected to be limited. Indeed, in some cases the mRNA is detected throughout preimplantation development, but the protein is only observed from a certain preimplantation stage onward (Vinot et al. 2005); or the mRNA is degraded soon after fertilization, while the protein persists through the blastocyst stage (Coonrod et al. 2006; Li et al. 2008; Ohsugi et al. 2008). Unfortunately, conventional tools for protein analysis such as antibodies (immunofluorescence, immunocytochemistry, western blotting) do not scale well to genome-wide investigations.

Large-scale qualitative and quantitative proteomic technologies have matured over the past two decades. In particular, direct measurement of proteins using mass spectrometry (MS) holds great promise as a complement to transcriptomics. Still, current high-throughput protein quantification methods are less sensitive than those for mRNA, and until a few years ago, the difficulty in obtaining sufficient amounts of input material had made the analysis of the mammalian oocyte and embryo proteomes utilizing MS effectively prohibitive. Prior to this study, 7,000 mouse oocytes/zygotes per sample had been required to identify ~3,000 proteins up to the 1-cell stage (Wang et al. 2010), while 3,000 mouse blastocysts per sample had been necessary to determine ~2,500 proteins (Fu et al. 2014). Very recently, increasing the input amount to 4,000-8,000 embryos per sample proved sufficient to distinguish ~5,000 proteins during mouse development from 1-cell stage to blastocyst (Gao et al. 2017). Bovine oocytes and embryos are larger, and by analyzing 100 of them (Deutsch et al. 2014; Demant et al. 2015), ~1,000 to 1,500 proteins were detected. While based on different MS technologies, all these studies shared high numbers of oocytes or embryos, far greater than the single *Xenopus* or the few *Drosophila* oocytes that were sufficient to detect ~5,000 proteins (Kronja et al. 2014; Smits et al. 2014; Sun et al. 2014; Casas-Vila et al. 2017). The input amount and the size of the detected proteome correlate, and mammalian oocytes and embryos are small. Thus, without mass-killing oocyte donors or mass-producing oocytes from stem cells *in vitro* (Hikabe et al. 2016), we need to achieve more with less.

We were confronted with two major challenges constraining the systematic large-scale protein analysis of the mouse embryo, namely: 1) the requirement for astounding numbers of oocytes or embryos, and 2) the lack of information on dataset complexity and completeness. We combined high-throughput liquid chromatography-tandem mass spectrometry (LC-MS/MS) with mRNA sequencing to generate datasets encompassing seven stages of mouse development spanning from the oocyte to the blastocyst. We anticipate that this resource will be key to gaining a greater understanding of the oocyte to embryo transition, and provide two examples of its varied applications: we describe (1) how to query the ‘rule’ of weak transcript/protein correlation in order to expose exceptions to the rule; and (2) how to expand the list markers in order to follow the oocyte-to-embryo transition.

Our dataset enriches the status of the mouse as a model system in developmental biology with the protein dimension, enabling a better understanding of the gene expression cascade that leads to the phenotype.

## Results

### Ultrastructural data underscore the relevance of a direct examination of the embryonic proteome

To systematically investigate the relationship between the proteome and the transcriptome in the developing mouse, we chose the paradigm of recovering fertilized oocytes *in vivo* after ovarian stimulation and culturing them *in vitro* in KSOM(aa) medium under 5% CO_2_ in air (see Methods). This made it possible to continuously monitor the progression of the embryos, to identify and collect stages more precisely, and to allay concerns over the quality of embryos developing inside a stimulated genital tract (Ertzeid and Storeng 2001; Van der Auwera and D’Hooghe 2001). In a separate group of embryos used to test for developmental quality, 89.5% (N=258) of the fertilized oocytes developed to blastocyst and, of these, 42.3% (N=104) progressed to term (embryo transfer). Typical features of early mouse development, including changes in endoplasmic reticulum (ER) architecture (Cech and Sedlackova 1983) and in ribosome morphology (van Blerkom and Brockway 1975; Bachvarova et al. 1981; Piko and Clegg 1982) were recapitulated, supporting the use of our *in vitro* system to yield embryos that are representative of normal development. In particular, we noted that hexagonal-shaped free ribosomes enabling efficient protein synthesis (van Blerkom and Brockway 1975; Bachvarova et al. 1981; Piko and Clegg 1982) are rare prior to the morula stage (see Figure 1A). Nevertheless, and in agreement with previous studies (Kidder and McLachlin 1985; Latham et al. 1991), developmental progression was impeded when cycloheximide, an inhibitor of protein synthesis, was added to the culture medium (see Figure 1B), indicating that protein synthesis is essential for further development of the early embryo.

**Figure 1.**
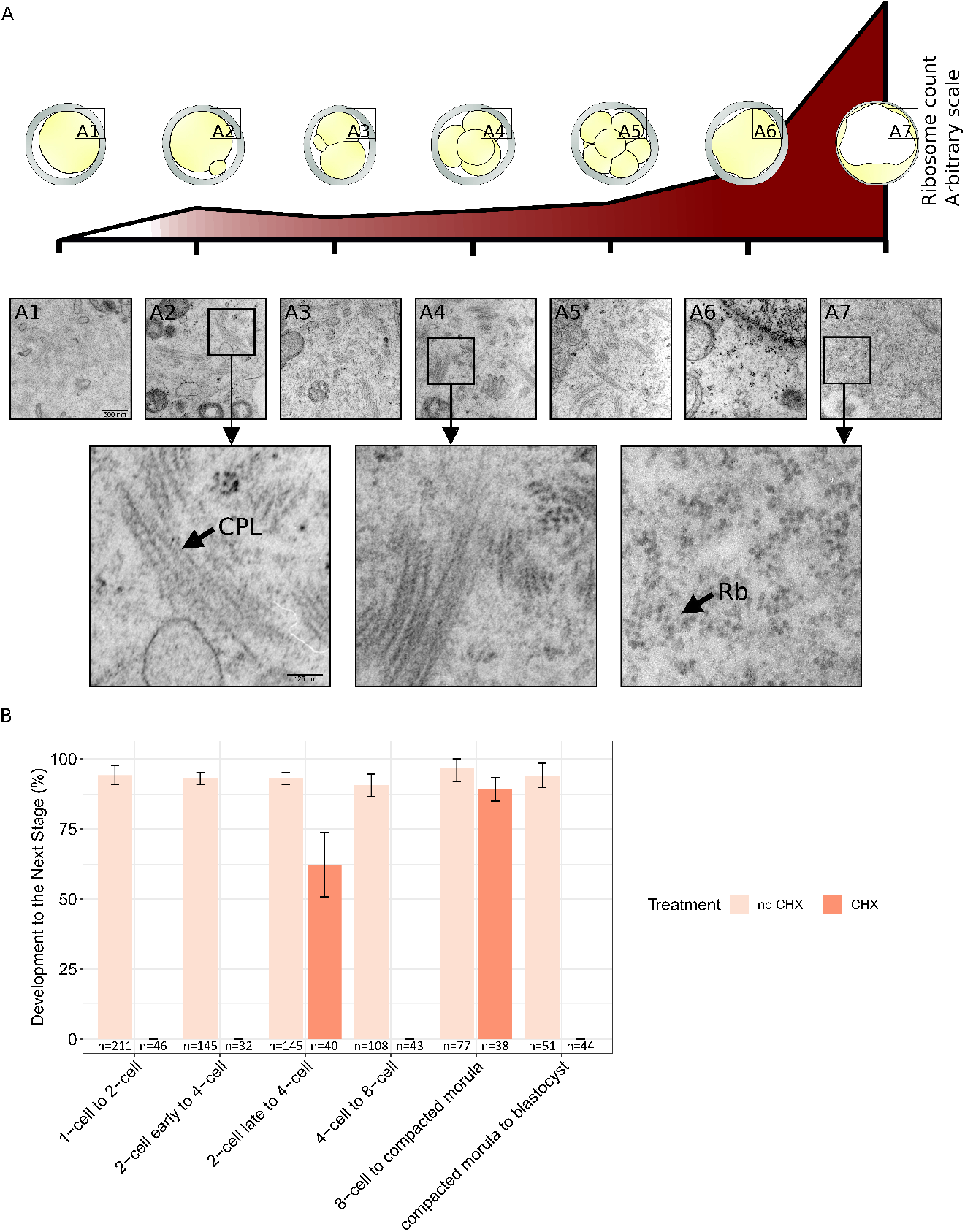
In vivo-fertilized, in vitro-cultured mouse oocytes as a source of embryonic material for proteomic analysis. **(A)** Oocytes and developmental stages were examined in ultrastructure. The density of hexagonal-shaped free ribosomes increases over time during preimplantation development (estimates based on 3 sections from three different embryos of each stage). Micrographs of cytoplasmic lattices (black arrow, “CPL”) and free ribosomes (black arrow, “Rb”) are shown. **(B)** The treatment of embryos with cycloheximide (CHX) documents that protein synthesis is necessary for developmental progression to the next stage. For each stage transition, height of the bars denote the percentage of embryos developing to stage without adding CHX; orange histograms denote the same percentage (if any) after treatment with CHX. The numbers under the bars indicate the total number of embryos examined.

Together, these data suggest that the impact of transcriptional changes on the proteome may be small, calling for a direct examination of the embryonic proteome.

### A high-quality proteome of mouse oocytes and preimplantation embryos to a depth of 6,550 proteins

For the proteome analysis we collected and processed a total of ~12,600 oocytes or embryos, in three biological replicates of ~600 oocytes/embryos per developmental stage: unfertilized oocytes, fertilized oocytes with pronuclei, and preimplantation embryos at the 2-, 4-, 8-cell, advanced morula and blastocyst stages (see Methods). The detected proteome comprised 6,550 proteins. Among these, 5,217 proteins were detected in at least two replicates of one or more developmental stages, and 1,709 proteins were detected in all replicates of all developmental stages. Protein abundance measurements (L/H ratios, see Methods) were highly reproducible, with minimum Spearman’s rank correlation coefficients between replicates in the range of 0.67 to 0.76 (for the oocyte and 2-cell stage, respectively, see Supplemental Fig. S1). Compared to the theoretical proteome (see Supplemental Methods), the 6,550 detected proteins are mainly involved in RNA processing, organelle organization, intracellular transport and cellular metabolism. Although these processes are not exclusive to preimplantation development, they are consistent with the nature of embryonic cleavage as a phase of development during which biomass is not produced *de novo* but rather reorganized.

To date, the scientific literature offers no estimate of the size of the mouse preimplantation proteome. While the number of distinct proteins in a cell line can be estimated by performing replicates at will, until the number of distinct proteins in the aggregated replicates approaches saturation, this is not feasible with mammalian oocytes and embryos. Therefore, we adopted a metabioinformatics approach. First, for each of 233 known mammalian protein complexes (based on (Ori et al. 2016), see Supplemental Methods), we computed the fraction of its members that are present in our dataset. Since all its members are required for the function of a complex, undetected members hint at a technical limitation rather than genuine biological absence. The overall median for the fractions of complex members detected in at least one replicate was 0.80, and ranged from 0.75 to 0.80, depending on the developmental stage (see Supplemental Fig. S2). Furthermore, we investigated how these fractions depend on the preferential cellular localization of the complexes (see Supplemental Methods), obtaining medians from 0.45 (cytoskeleton in the blastocyst) to 1.00 (endosome at all developmental stages, see Supplemental Fig. S3). Similarly, among individual replicates we found overall medians of 0.44, 0.67 and 0.78 (see Supplemental Fig. S4, S5 and S6). Second, we aggregated the replicates of each developmental stage and studied the cumulative number of distinct proteins as a function of the number of aggregated replicates. Random replicate aggregation resulted in an apparent saturation at ~5,500 to 6,000 proteins, with each replicate leading to only a small increase in the number of proteins detected (see Supplemental Fig. S7). Likewise, aggregating samples from different developmental stages in random order resulted in rapid saturation at ~6,250 proteins, such that the first four samples contribute the vast majority of distinct proteins (see Supplemental Fig. S8). The same is valid for different groups of proteins expected to be present in very different concentration ranges, such as transcription factors, enzymes and structural proteins (see Supplemental Methods and Figs. S9, S10 and S11). Third, we directly compared our dataset to a very recently published dataset in which 4,830 different proteins were identified in at least one of two replicates from six developmental stages (1-cell to blastocyst, (Gao et al. 2017)). We found that 4,028 (83%) of these proteins are contained in a reduced version of our dataset comprising the same six developmental stages (see Supplemental Fig. S12). In particular, 378 (81%) of the structural proteins, 1,180 (89%) of the enzymes and 180 (88%) of the transcription factors in this dataset are contained in ours. Moreover, our reduced dataset contains an additional 2,369 proteins (not present in the dataset of (Gao et al. 2017)), among which are 172 structural proteins, 537 enzymes and 188 transcription factors. Together, these findings suggest that we have achieved a high coverage – of up to 80% – of the mouse preimplantation proteome.

In summary, the dataset described here is of high quality and can be useful for in-depth investigation of early mammalian development and hypothesis generation or testing.

### The dynamics of protein expression orchestrating preimplantation development is complex

As described numerous times on the mRNA level, fertilization is followed by extensive gene expression reprogramming. Nevertheless, the impact of transcriptional changes on the proteome is uncertain. Thus, it has been hypothesized that once activated, a gene continues to be transcribed during later developmental stages, resulting in product accumulation (Kidder 1992) that extends into the proteins. On the other hand, early protein studies of the mouse embryo based on radioactive gel electrophoresis support the hypothesis that protein expression occurs in phases (Levinson et al. 1978). To identify proteins whose expression significantly fluctuates as a function of the developmental stage, we subjected the 5,217 proteins detected in at least two replicates of at least one developmental stage to an analysis of variance (ANOVA, see Methods). This revealed a total of 1,290 (25%) differentially expressed proteins (P-value ≤ 0.05). Among these, 905 proteins exhibited fold changes ≥ 2 or ≤ 0.5 between any two developmental stages. In particular, 488 proteins showed fold changes ≥ 2 or ≤ 0.5 between consecutive developmental stages (see Figure 2A); most (253) of these exclusively during the transition from the morula to the blastocyst (see Supplemental Fig. S13). Compared to the detected proteome, the 488 proteins were associated with small molecule and carboxylic acid/carbohydrate metabolism, enzymatic activity; and (extracellular) exosome production (FDR ≤ 0.05, see Supplemental Methods and Table S1). These terms are consistent with a sequence of landmark events in mouse preimplantation development, such as the enzymatic transition from metabolic usage of pyruvate to usage of glucose (Dumollard et al. 2007), and the paracrine communication between embryos (Saadeldin et al. 2014) as well as between the embryos and the maternal genital tract (Giacomini et al. 2017).

**Figure 2.**
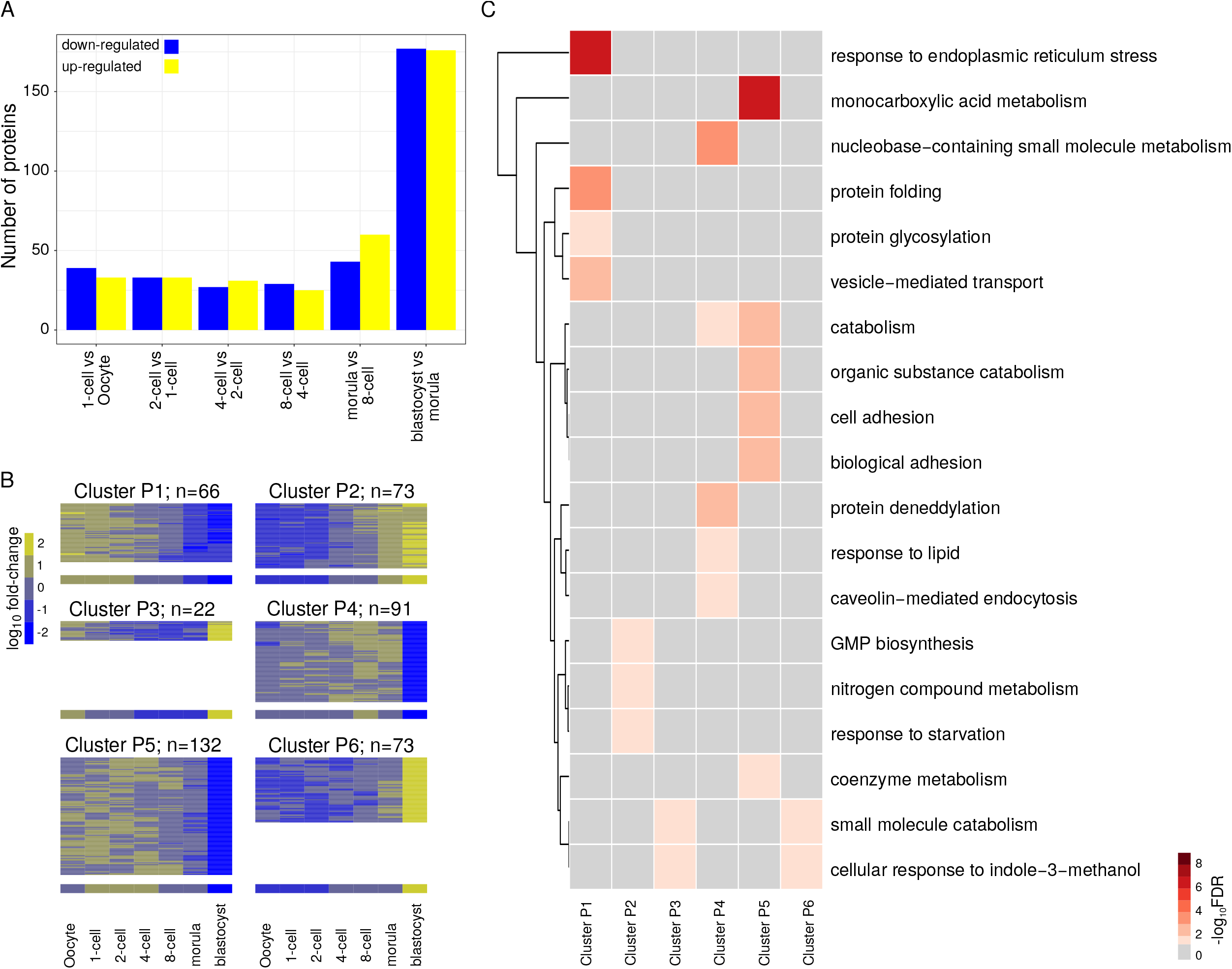
Differentially expressed proteins across preimplantation development and their functional enrichment. **(A)** Number of differentially expressed proteins between pairs of consecutive developmental stages (fold-change ≥2 or ≤ 0.5 between any two developmental stages, P-value ≤ 0.05 from ANOVA). **(B)** Expression profile of protein clusters. The heatmaps show fold-changes relative to the oocyte scaled using the z-score transformation. The height of the heatmaps is proportional to the number of proteins in each cluster, which is also indicated. The median fold-change across all cluster members for each developmental stage is represented below the heatmaps. (C) Annotation of protein clusters. Gene Ontology (GO) terms associated (FDR ≥ 0.05) with each cluster were summarized with REVIGO (Supek et al. 2011). REVIGO “representatives” for the individual GO terms are listed on the right of the heatmap. Statistical significance (sum of the -log_10_ FDR of the individual GO terms) for the annotation of each of the clusters is represented using a color gradient. Non-significant associations are represented in gray. Details are presented in Supplemental Table S2.

Fuzzy clustering of the 772 proteins detected in at least two replicates of each developmental stage that were differentially expressed (P-value ≤ 0.05) and showed a fold change ≥ 2 or ≤ 0.5 between any two developmental stages revealed six clusters (see Supplemental Methods and Figures 2B and C). The two largest clusters (P5 and P4) comprise proteins whose expression decreases sharply between the morula and blastocyst stages and, compared to the detected proteome, are primarily enriched in monocarboxylic acid metabolism (P5), and nucleobase-containing small molecule metabolism (P4, see Figure 2C and Supplemental Table S2). Clusters P5 and P4 are approximately mirror images of clusters P6 and P3, respectively. Nevertheless, the proteins in clusters P6 and P3 have their own functional profiles; thus, both clusters are connected to small molecule catabolism and cellular response to indole-3-methanol. The two remaining clusters (P1 and P2) are mirror images of each other and comprise proteins that steadily decrease or increase towards the blastocyst. Proteins in cluster P1 are mainly associated with response to endoplasmic reticulum stress and protein folding, whereas those in cluster P2 are related to GMP biosynthesis, nitrogen compound metabolism and response to starvation.

To build more confidence in the observed protein profiles, we independently validated our proteomics measurements using immunofluorescence assays. Since validation is impracticable for all proteins, we selected three proteins with different expression profiles. Among the proteins that are present throughout development (albeit more abundant at the beginning) we selected Ddx6, which is associated with processing bodies (P-bodies) involved in the storage and degradation of mRNAs (Hu et al. 2010). Among the proteins that are detected in oocytes and early stages but become undetected later on, we selected Rc3h1, also known as roquin, which is an element of a post-transcriptional repression pathway and whose mutation leads to the *sanroque* phenotype (Vinuesa et al. 2005). Among the proteins that are not detected in oocytes and early stages but become detected later on, we selected Alppl2, known for its role in the placenta and expressed in the trophectoderm of the preimplantation embryo (Johnson et al. 1977; Hahnel et al. 1990; Bai et al. 2012). The immunofluorescence profiles of Ddx6, Rc3h1 and Alppl2 matched the corresponding proteomics profiles (see Supplemental Fig. S14). Moreover, to extend the analysis we collected and curated enzymatic/immunofluorescence data from the literature and/or obtained in the past by our own laboratory on 33 proteins (37 sets of measurements across multiple developmental stages, see Supplemental Table S3). Specifically, we quantified the similarity between the expression profiles as determined by enzymatic/immunofluorescence assays and our proteomics pipeline by computing the Spearman’s rank correlation. We observed strong correlations (Spearman’s rank correlation coefficients between 0.6 and 0.79) for seven proteins (and seven sets of measurements) and very strong correlations (Spearman’s rank correlation coefficients between 0.8 and 1.00) for five proteins (seven sets of measurements). The results are significant compared to the random expectation (empirical P-value < 0.006, see Methods).

Taken together, these results reveal systematic changes of the proteome of the embryo as it develops. Furthermore, these changes are complex and unlikely to reflect a mere alternative between monotonic accumulation and stage-specific expression (Levinson et al. 1978; Kidder 1992).

### Changes in protein abundances become more prominent as development progresses, and so does the concordance with changes in transcript expression values

Previous transcriptome-based studies of mouse embryonic development have shown that the transcriptomes of oocytes and early embryos can be clearly divided into two groups: prior and after the 2-cell stage (Hamatani et al. 2004; Wang et al. 2004; Zeng et al. 2004). However, the most conspicuous morphological changes during preimplantation development – compaction and cavitation – occur well after the 2-cell stage, in the morula (Hamatani et al. 2004). To directly compare the oocyte-to-embryo transition on the protein and mRNA levels, we generated our own transcriptome using RNA-seq. For this purpose, we collected and processed a total of 3,424 oocytes or embryos in two biological replicates of 214 oocytes/embryos per developmental stage (see Methods). Anticipating major differences between the early and late 2-cell stage, we considered these separately. We identified a total of 20,535 protein-coding transcripts with at least one read count in any of the samples.

As expected, principal component analysis (PCA) of the expression values of the transcripts showed that developmental stages can be distinguished based on their transcriptomes and that most of the variance in the data is contributed by changes at early developmental stages, (see Figure 3A). PCA performed on the abundances of the cognate proteins also clearly distinguished the developmental stages based on their proteomes, albeit with most of the variance in the data being contributed by changes between the morula and the blastocyst. Indeed, the progression from the 4-cell to the blastocyst embryo aligned almost perfectly with the increase of the first principal component (PC) and explained 31.1% of the variance, while the progression from the oocyte to the 4-cell stage aligned with the decrease of the second PC and explained only 11.8% of the variance (see Figure 3B). These findings indicate that in contrast to the transcriptome, in the proteome the oocyte-to-embryo transition is less connected to the 2-cell stage, with the protein expression signature of the blastocyst being particularly further apart from those of the other developmental stages considered. This is in agreement with the establishment of two blastocyst cell populations that differ radically in their metabolic and cell cycle parameters: polarized external cells (the future trophectoderm) and apolar internal cells (the future inner cell mass) (MacQueen and Johnson 1983; Houghton 2006).

**Figure 3.**
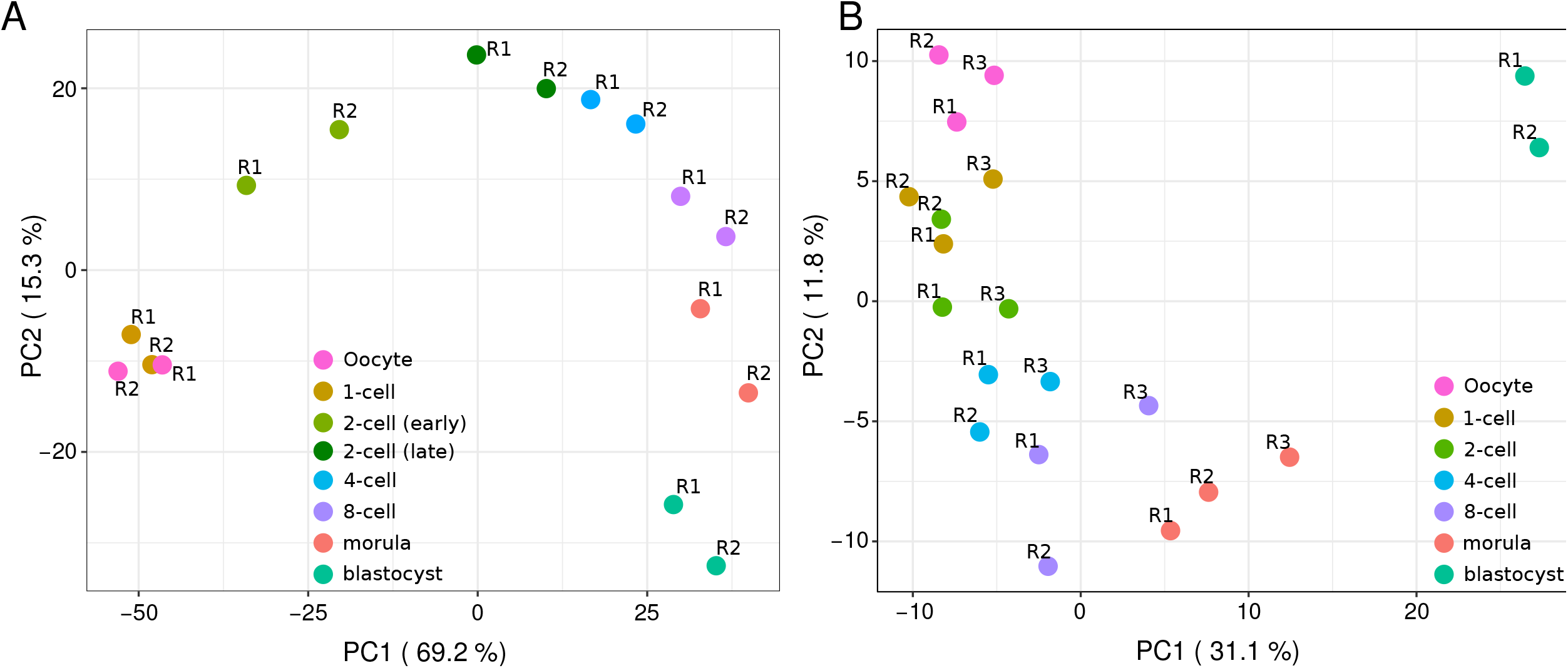
The proteome and the transcriptome develop differently in time. **(A)** Principal component analysis (PCA) of the expression values of the detected transcripts that are the cognates of the proteins detected in all replicates at all developmental stages (see B). The first two PCs are shown, with sample points colored by developmental stage. **(B)** Principal-component analysis of the (log_2_) L/H ratios of the 1,709 proteins detected in all replicates of all developmental stages.

To explore the relationship between the transcriptome and the proteome in the course of preimplanatation development, we first investigated the correlation between the (log_2_) fold-changes in protein abundances and the expression values of the cognate transcripts relative to the oocyte. We found a strikingly weak correlation, with Spearman’s rank correlation coefficients in the range of -0.06 (1-cell early versus 2-cell) to 0.41 (morula versus blastocyst, see Supplemental Fig. S15), confirming important differences between the protein and transcript expression profiles of the developing embryo.

Next, we divided the proteins into two disjoint groups according to the direction of change in expression of their cognate transcripts relative to the oocyte (see Figure 4A). More precisely, given developmental stages *S_i_* and *S_j_*, we separated the proteins into two groups: i) those with transcripts up-regulated at *S_i_*; and ii) those with transcript down-regulated at *S_i_*. Then, for each of the two groups of proteins, we estimated the probability of observing a certain protein (log_2_) fold-change at *S_j_* relative to the oocyte (see Figure 4B and C and Supplementary Methods). For any (log_2_) fold-change *x*, if the protein expression changes at *S_j_* reflect the transcript expression changes at *S_i_*, the probability of observing a protein with a (log2) fold-change of *x* or less at *S_i_* is expected to be greater for those proteins whose transcripts are down-regulated than for those whose transcripts are up-regulated at *S_i_*. Hence, we quantified the concordance between protein and transcript expression changes by measuring the difference between the areas bounded by the two implicit cumulative distribution functions (CDFs, see Supplementary Methods). This analysis revealed little or no concordance between protein and transcript expression changes at early developmental stages (see Figure 4D and Supplemental Fig. S16). The concordance, however, increased towards later developmental stages, with expression changes at the morula and blastocyst stages exhibiting the overall highest concordances. Moreover, the concordance for the transcript expression changes at the 4-cell, 8-cell and morula stages was highest for the protein changes at the blastocyst stage, and higher than that between transcript expression changes at the blastocyst stage and protein changes at the blastocyst stage (see Figure 4D).

**Figure 4.**
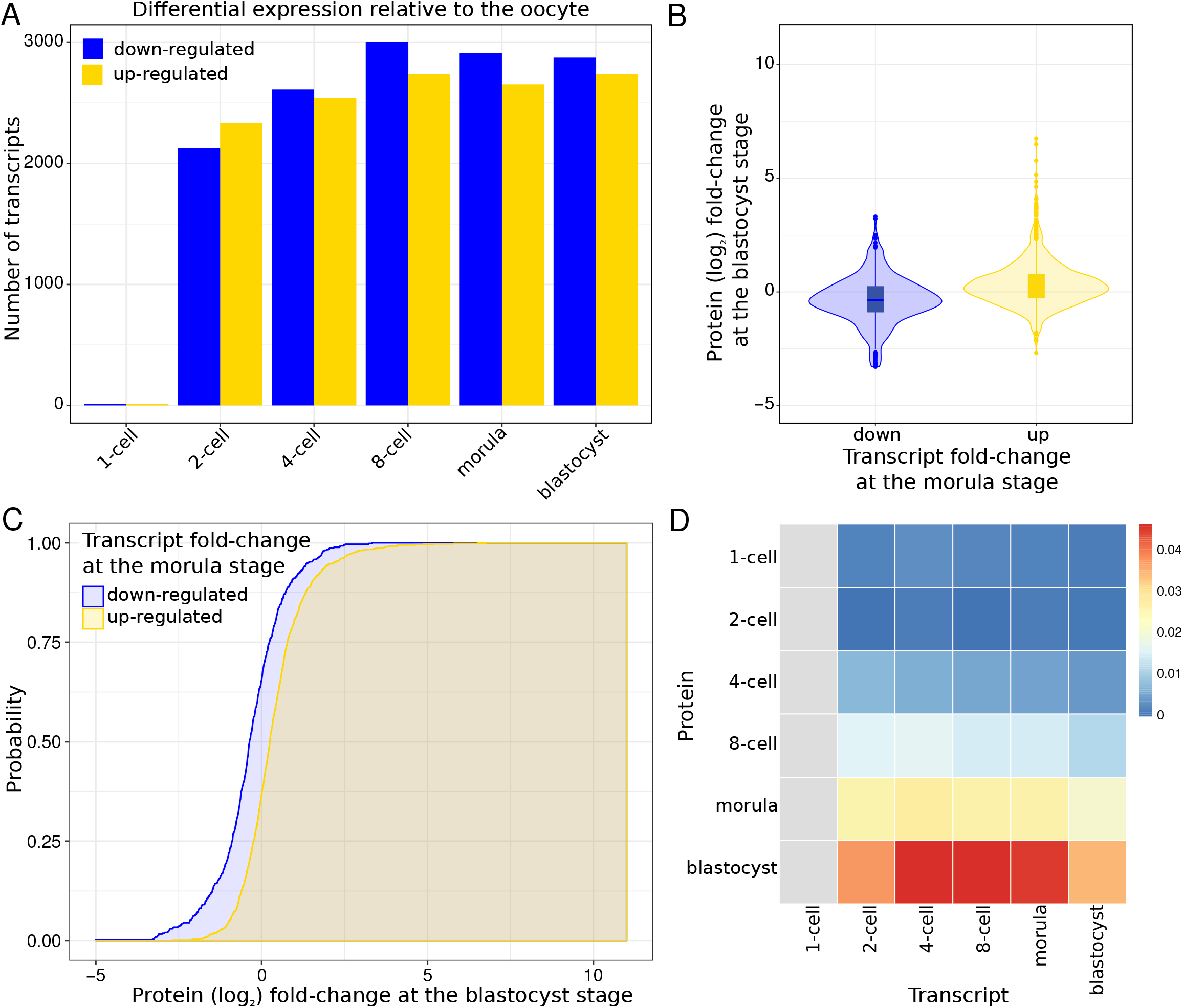
Changes in the transcriptome are reflected at the proteome level from the morula stage onwards. **(A)** Number of differentially expressed transcripts at each developmental stage relative to the oocyte (FDR≤1×10^-5^ and fold-change ≥2 or ≤ 0.5 between any two developmental stages). **(B)** Violin plot showing the distribution of protein (log_2_) fold-changes at the blastocyst stage relative to the oocyte for proteins whose cognate transcripts are down-(blue) or up-(yellow) regulated at the morula stage relative to the oocyte. **(C)** Cumulative density functions (CDFs) of protein (log_2_) fold-changes at the blastocyst stage relative to the oocyte for proteins whose cognate transcripts are down-(blue) or up-(yellow) regulated at the morula stage relative to the oocyte. 1,617 proteins and their cognate transcripts were used to estimate the CDF: 5,565 transcripts were found differentially expressed between the morula and the oocyte; 1,617 of the cognate proteins were detected in both the blastocyst and the oocyte. The CDF for the proteins whose transcripts are up-regulated is shifted to the right compared to the CDF for the proteins whose transcripts are down-regulated. We used the difference between the two areas bounded by the CDFs (shaded) to quantify the shift and, thereby, the impact of the transcript expression changes at the morula stage on the protein expression changes at the blastocyst stage. **(D)** Heatmap representing the difference in the area of the two CDFs for all pairs of stages. The x-axis of the grid corresponds to transcripts; the y-axis corresponds to proteins. The colors indicate the differences between the two areas bounded by the corresponding CDFs normalized using the entire protein fold-change range. Red indicates a large area, and hence a considerable shift between the distributions, while blue indicates the opposite. Gray indicates the pairs of stages for which we did not estimate the CDFs because less than 25 proteins/transcripts were detected and/or found differentially expressed, respectively.

Altogether, these results are in agreement with the increase in the density of free ribosomes that enable efficient protein synthesis only starting at the morula stage (see Figure 1A). Despite some *de novo* transcript synthesis beginning at the 1-cell stage, the lack of a conventional translation machinery (i.e., the lack of free ribosomes) prevents transcripts from being robustly translated until the morula stage. Furthermore, despite the steep increase of the free ribosomes, a delay between transcription and translation is still evident at the blastocyst stage. Overall, the majority of the proteins do not match the previously described (Ko et al. 2000) stage-specific groups of transcripts that support a ‘hit and run cascade’ model for early embryonic development. Instead, our results document only a moderate amount of change in the proteome, suggesting a steady basal translation of transcripts into proteins, and a role for subcellular compartmentalization and storage in order to make the proteins available when and where required.

### Exceptions to the rule of weak transcript-protein correlation define a special class of genes with distinct developmental functions

To exemplify how our dataset can be analyzed to better understand the relationship between the transcriptome and the proteome during preimplantation development, we applied fuzzy clustering to the transcripts of the 772 proteins that we had clustered before, and compared the transcript and protein clusters. Specifically, we clustered the (log_2_) fold-changes of the transcripts relative to the oocyte, and found seven clusters (see Supplemental Methods, Fig. S17 and Table S4), which, in contrast to the protein clusters, are often characterized by expression profiles with evident inflection points either at the early or late 2-cell-embryo stage. Next, to quantify the similarity between the protein and transcript clusters, we computed the Pearson’s correlation coefficient ( ) between their expression profiles in a pairwise manner (see Methods). Out of a total of 42, 14 pairs had a Pearson’s correlation coefficient greater than or equal to 0.5, indicating high similarity (see Figure 5A). In addition, we assessed the overlap between the members of all pairs of protein and transcript clusters and found that only ten shared more proteins/transcripts than expected by chance (P-value ≤ 0.05, one-sided Fisher’s exact test, see Figure 5B). The overlap was particularly high among pairs of protein and transcript clusters with similar expression profiles (*r* ≥ 0.5), with an odds-ratio of 7.8 (P-value=0.008, one-sided Fisher’s exact test), highlighting the fact that, despite the little overall concordance between protein and transcript expression changes, the expression of some proteins indeed mirrors that of their cognate transcripts. Compared to the 772 differentially expressed proteins and their cognate transcripts considered for clustering, the 146 genes that overlap among the pairs of clusters with similar expression profiles were enriched in positive regulation of secretion, reflecting the increasing role of the embryo-derived ‘secretome’ as development progresses in preparing the ground for the molecular dialogue between the embryo and the maternal endometrium (Giacomini et al. 2017).

**Figure 5.**
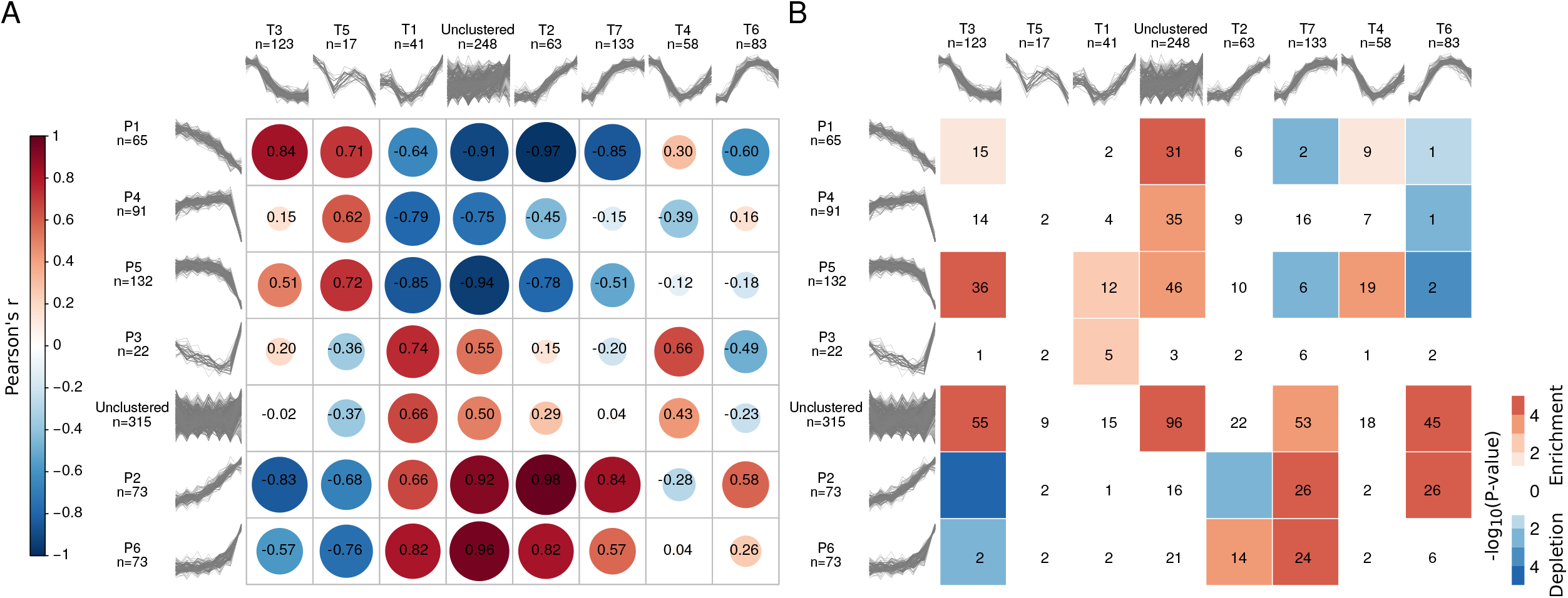
Comparison between protein and transcript clusters. **(A)** Pearson correlation coefficients (in black) computed between the medians of the expression profiles of pairs of protein and transcript clusters. The correlation matrix based on the Pearson correlation coefficients between the median (log2) fold changes across the members of protein and transcript clusters was visualized using the R corrplot package (Wei and Simko 2017). The size and color of the circles are both indicators of the magnitude and sign of the correlation. The matrix was reordered based on hierarchical clustering using the complete linkage method. **(B)** Significance of the overlap (-log10 P-value calculated using Fisher’s Exact test) between the members of protein (rows) and transcript (columns) clusters. Significant enrichments are depicted in red and depletions in blue. The numbers indicate the size of the overlap; overlaps of size zero are not indicated.

For the purpose of identifying and characterizing the genes with either strongly correlated or anticorrelated protein and transcript expression profiles, we perused the correlation between the expression profiles across the developmental series of the aforementioned 772 differentially expressed proteins and their cognate transcripts. Despite the expected weak overall correlation (with a median Spearman’s rank correlation coefficient across all genes of 0.18), we observed that the distribution of Spearman’s rank correlation coefficients was relatively broad (see Supplemental Fig. S18). To enhance confidence in the observed profiles, we independently validated our proteomics and transcriptomics data using immunofluorescence and TaqMan (qPCR) respectively (see Methods). We selected three genes among those exhibiting strong negative correlations during preimplantation development and particularly from the 1- to the 4-cell stage: Pdia3, Top1, and DNAjb11. These genes are particularly appropriate in the context of our study, because mutations of Pdia3, Top1 and Dnajb11 interfere with development and prove lethal in homozygosis (Morham et al. 1996; Francisco et al. 2010; Li et al. 2014a; Wang et al. 2014). The results of the TaqMan assay for Pdia3, Top1 and Dnajb11 correlated positively with those of RNA-seq, as did the results of the immunofluorescence with those of LC-MS/MS (see Supplemental Figure S19), confirming the existence of genes with strongly anticorrelated protein and transcript expression profiles.

Finally, with the comfort of the validation data, we moved on to analyze the features of the genes at the extremes of the distribution of Spearman’s rank correlation coefficients. Indeed, 7% of the proteins and transcripts exhibited very strong positive correlations (≥ 0.8) and 3% showed very strong negative correlations (≤ -0.8). Among the former are genes involved in ubiquitin metabolism and ubiquitination (Dcun1d5, Uspx9, Dcaf8, Gabarapl2, Rnf114, Stt3b, Ube2g1), signal transduction (Arhgap12, Gna13, Pdpk1), synthesis and modification of DNA (Ctc1, Rrm2, Hmces), splicing and storage of mRNA, including translational initiation (Paip1, Rbm8a, C1qbp, Igf2bp3, Igf2bp2, Xab2, Nhp2). Among the latter are genes involved in membrane vesicle trafficking (Dynlrb1, Epn2, Vta1, Napa, Eea1), chaperoning (Hypk, Fkbp2), and protein glycosylation in association with ribosome binding (Rpn1, Rpn2). Known genes with established roles in development are found in both groups (e.g., Rrm2 (Yu et al. 2016); Igf2bp2 (Dai et al. 2015); Igf2bp3 (Li et al. 2014b); Epn2 (Chen et al. 2009)). Overall, the proteins with very strong positive correlations are implicated in dynamic processes, while those with very strong negative correlations represent maintenance systems, with a convergence on signaling.

Thus, the release and uptake of vesicles supported by the anticorrelated genes is one way to modulate the concentration of signaling molecules supported by the highly correlated genes, as exemplified by the case of Epn2 (Chen et al. 2009).

### Proteomic profiles suggest new markers to better follow the oocyte-to-embryo transition

To show how our dataset can be applied to the identification of new candidate developmental markers, broadening the options offered by morphology/morphokinetics or metabolic markers secreted into the culture medium, we examined the molecular basis of morphological staging. As an illustration, we uncovered new candidate markers to follow the oocyte-to-embryo transition, and thus compared the proteomes of early (oocyte, 1- and 2-cell embryos) and late (4-cell to blastocyst embryos) developmental stages. In particular, we trained and tested linear discriminant analysis (LDA) classifiers. Our results show that protein expression can be used to perfectly separate between early and late developmental stages, with an area under the Receiver Operator Characteristic (ROC) curve of 1.00 (see Supplemental Methods and Fig. S20). Samples from the 4-cell stage embryos were close to the decision boundary of the classifier, indicating at this stage the coexistence of features from both previous and later stages, and characterizing the 4-cell stage as a transitional stage. Further, we inferred 20 potential markers for early and late developmental stages by ranking the proteins according to their relevance for the classification (see Supplemental Methods and Figure 6A). These proteins include enzyme modulators, hydrolases and ligases (see Figure 6B). In particular, Ddx6 is an RNA helicase that has been found in P-bodies (Decker and Parker 2012) and is involved in translation repression and in 2-cell stage embryonic arrest (Hu et al. 2010). Moreover, some of these proteins (e.g., Ppm1a and Wtap) are mediators of TGF-β and Wnt signaling (Lin et al. 2006; Wu et al. 2016). This finding is compatible with the aforementioned overrepresentation of ‘exosome production’ among differentially expressed proteins, since signaling pathways rely in part on exosome-mediated mobilization. Interestingly, five of the 20 markers (Calr, Hyou1, Pdia3, Pdia4 and Txndc5) are involved in the protein processing in endoplasmic reticulum (ER) pathway (KEGG identifier mmu04141 (Kanehisa and Goto 2000; Kanehisa et al. 2016; Kanehisa et al. 2017), odds-ratio = 6.7, P-value = 0.002, Fisher’s Exact test), enlightening the molecular basis of the changes in ER architecture that take place during the transition from oocyte to embryo (Kim et al. 2014) and that are concomitant to the increase in protein synthesis and folding after EGA (Michalak and Gye 2015). These twenty marker proteins constitute good candidates for further molecular studies of mammalian preimplantation development.

**Figure 6.**
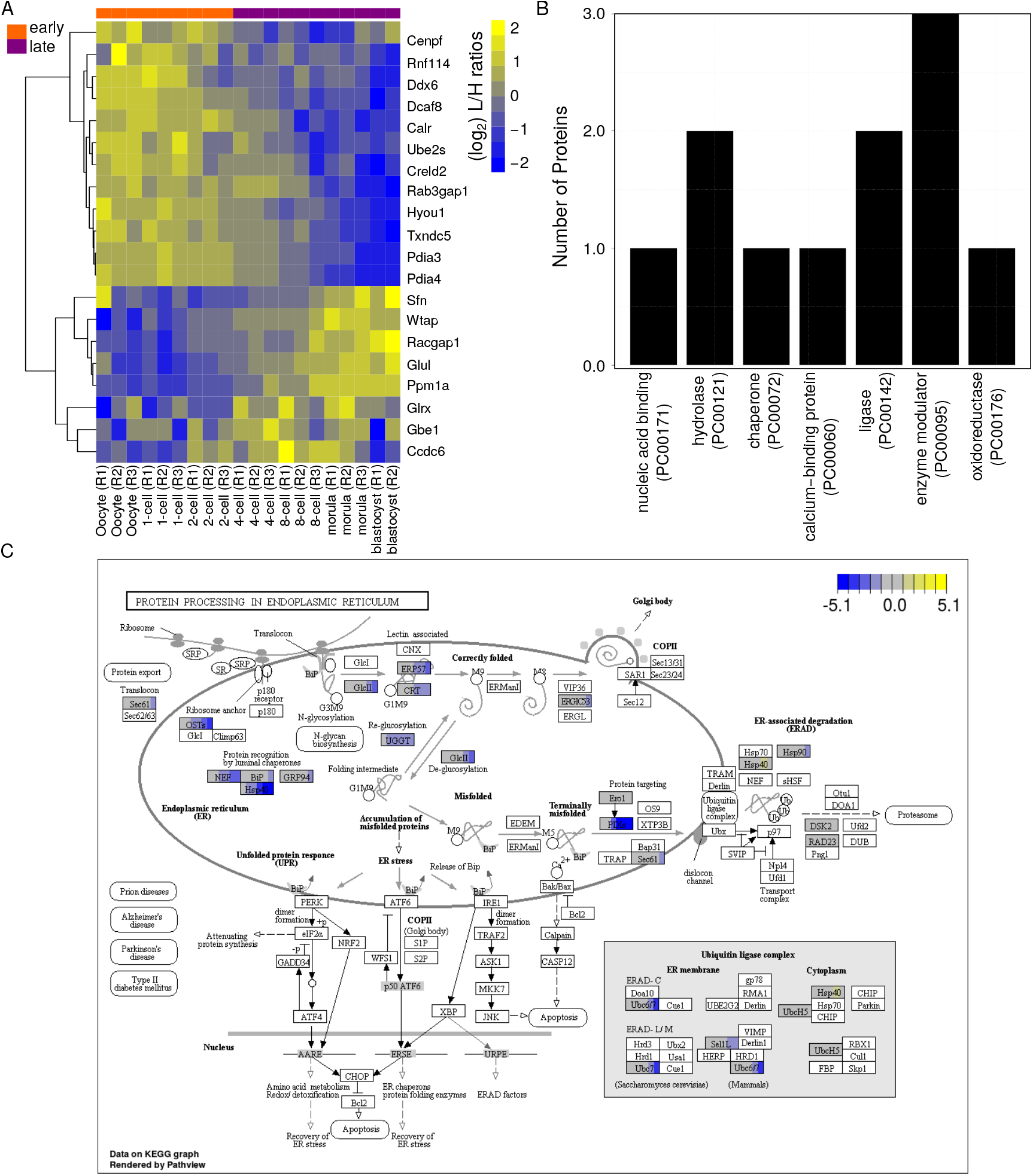
Classification of early and late preimplantation developmental stages based on protein abundances using Linear Discriminant Analysis (LDA). **(A)** Heatmap of (log_2_) L/H ratios for 20 candidate protein markers for distinguishing between early and late preimplantation developmental stages. The 20 samples are sorted chronologically according to developmental stage and replicate number. Row clustering was performed with a Pearson correlation-based distance using the complete linkage method. The package pheatmap in R was used for visualization (Kolde 2016). **(B)** PANTHER protein classification available for 11 of the 20 candidate proteins markers (see A). **(C)** Pathway analysis of genes differentially expressed in the murine “protein processing in endoplasmic reticulum” KEGG pathway (mmu04141). The boxes representing the proteins/genes are uniformly divided by the number of developmental stages. Replicate averages are laid out chronologically from left to right across all developmental stages considered. The (log_2_) of the fold-change relative to the oocyte is indicated in yellow (up-regulated) or blue (down-regulated). Only proteins/genes among the 764 that were found differentially expressed across the developmental series are colored. The “Pathview” R/Bioconductor package (Luo and Brouwer 2013) was used to generate the graphical representation of the pathway and to indicate which genes are differentially regulated.

## Discussion

Prior to this resource, molecular descriptions of mammalian development had typically been centered on transcripts, although only for a small fraction of the known protein-coding transcripts there was actual proof of the presence of the corresponding proteins in the mouse embryo. In this study, we used MS-based proteomics to generate a proteome dataset with three biological replicates for the preimplantation stages of mouse development, from the oocyte to the blastocyst. This proteome was compared to the cognate transcriptome generated by RNA-seq. With 6,550 detected proteins, ours is the largest developmental proteome of a mammalian species characterized to date, and yet substantially smaller than the number of 20,535 protein-coding transcripts found in the same samples. A similarly conceived, recently published study conducted with a different workflow (TMT instead of SILAC) revealed nearly 5,000 proteins despite the much higher amount of input material used (Gao et al. 2017). While neither of these datasets is complete, we found that our proteome coverage is in the order of magnitude of up to 80%. Clearly, most mRNAs are stored and only translated when needed, and MS-based proteomics of developmental stages is not solely a matter of input amount: it is largely a matter of sample preparation and preprocessing (e.g., prefractionating) and of the experimental procedures and equipment used.

Our main finding when taking the sole proteome into consideration is that the majority of detected proteins change only moderately in abundance during the development from oocyte to morula. Accordingly, we hypothesize that the oocyte-to-embryo transition may last until the morula stage, in contrast to the swifter transition at the transcriptome level, largely accomplished between 2-cell and 4-cell stage. The blastocyst’s proteome stands out as markedly different from the proteomes of the preblastocyst stages. This distinction is consistent with the formation of the first epithelium, the trophectoderm. Translation in the preimplantation embryo is limited by the availability of free ribosomes, which are the most active players of a cell’s translational machinery, but poorly represented in pre-morula-stage mouse embryos. This explains why an impaired translational machinery does not affect blastocyst formation, but causes blastocyst implantation failure in mice (Plaks et al. 2014).

Our main finding when comparing the protein abundance profiles with their cognate transcript profiles is that the projection of the proteome onto the developmental time axis differs from the prediction based on the transcriptome, with the correlation improving as development progresses. While most changes at the protein level explain the transition between the morula and the blastocyst, most changes at the mRNA level explain the transition between the oocyte and the 2- to 4-cell stage. Although the overall protein-mRNA correlation is weak, for a small subset (7%) of the detected proteome, the proteins and their cognate mRNAs have very similar profiles. Moreover, for another small subset (3%) of the detected proteome, the correlation is even negative, with protein levels increasing as transcript levels decrease. These cases may be explained, for example, by the packaging of RNA in granules, such as P-bodies (Hogan et al. 2008; Peshkin et al. 2015), whereby the mRNA broken free from these granules becomes available for both translation and degradation. Notably, we observed a decrease of the P-body protein Ddx6 from oocyte to blastocyst, which together with the increase in free ribosomes would explain the improving protein-mRNA correlation as development progresses. These covariates make the anti-correlated proteins virtually impossible to predict from their transcripts. From our data it is now clear that these anti-correlations are no exceptions, but manifestations of a non-negligible phenomenon in mouse development.

Two limitations of our study, apart from artifacts that may occur in our *in vitro* setting as well as in the *in vivo* situation (caused by the hormonal status of the genital tract; (Gao et al. 2017)), are the following. First, it is difficult to determine whether we failed to detect important proteins. However, our coverage estimates are in the order of magnitude of up to 80%, suggesting that the number of false negatives is bounded. Second, it is not known how the genotype of the gametes influences the composition of the developmental proteome. However, as reported by us (Pfeiffer et al. 2015), the proteomes of the oocytes of different inbred strains (129/Sv, C57Bl/6J, C3H/HeN, DBA/2J), while not identical, only differ in a minor proportion of detected proteins. A third limitation is that our ability to detect proteins in oocytes and embryos depends on the reference we used for SILAC. For example, trophectodermal markers seemed to be underrepresented in our dataset, although several of these proteins were also underrepresented in a study that did not use SILAC (Gao et al. 2017).

In summary, while there is still a long journey ahead until the proteome of mouse preimplantation development is exhaustively enumerated, our dataset constitutes a substantial contribution to closing the gap between ‘predicted’ phenotype (based on mRNA) and ‘actual’ phenotype (based on protein) of the mouse embryo, making an area of developmental processes now accessible to direct investigation as opposed to educated assumptions based on transcript levels. It is clear that the proteomic and transcriptomic analyses provide non-overlapping information except for a small subset of genes whose protein and mRNA profiles are highly correlated. The embryonic mouse proteome dataset described here will facilitate the study of mammalian development in at least two important ways. First, it will facilitate the molecular definition of embryo quality, which has a major impact on the course of gestation and yet is insufficiently accounted for on the molecular level. While morphological/morphokinetic markers commonly used to predict an embryo’s chances to develop can be subjective, our proteomic resource offers specific and measurable molecular candidates to complement the non-molecular markers. Thus, our LDA classifier was able to attain perfect separation between early and late developmental stages based solely on protein abundances. Second, since mammalian oocytes and embryos are produced in the gonads in comparatively small numbers (compared to e.g. Xenopus) and their availability can be subject to ethical and legal restrictions (e.g. in humans), knowing which gene products can be reliably predicted from mRNA has diagnostic value: these mRNA markers allow to make predictions that are backed by the proteins, and they do not require to consume the whole oocyte or embryo since cytoplasmic biopsies can be amplified for mRNA. For example, the cases of anti-correlation in which the mRNA is rapidly degraded after fertilization whereas the protein persist throughout the blastocyst stage, may be cases of candidate maternal genes. Beyond these examples, we believe that the range of applications of our resource is broad, depending on the personal interest of the user.

## Methods

### Ethics statement

This mouse study was performed in accordance with the recommendations of the Federation of Laboratory Animal Science Associations (FELASA) and with the ethical permit issued by the Landesamt fuer Natur, Umwelt und Verbraucherschutz (LANUV) of the state of North Rhine Westphalia, Germany (permit number: LANUV 81-02.04.2017.A432).

### Metaphase II oocyte collection

Metaphase II (MII) oocytes of B6C3F1 mice aged 8-10 weeks were collected from the oviductal ampullae after gonadotropin priming with 10 IU each PMSG and hCG injected 48 hours apart, as described (Wang et al. 2016; Casser et al. 2017).

### *In vivo* oocyte fertilization and *in vitro* embryo production

Gonadotropin-primed B6C3F1 females were mated to CD1 males (see Supplemental Fig. S21). Pronuclear oocytes were collected from oviductal ampullae at 10am on the day of the copulation plug. By 11am they had been freed of expanded cumulus cells in 50 U/mL hyaluronidase in HZCB medium, and placed in culture in 500 microliters KSOM(aa) medium (Ho et al. 1995) in 4-well plates (Nunc) under an atmosphere of 5% CO_2_ in air at 37 degrees Celsius. All embryos were staged carefully based on morphology and time spent in culture (beginning at 11am on the day of isolation from the oviduct).

### Transmission electron microscopy (TEM)

Mouse embryos were fixed 2h at room temperature in 2,5% glutaraldehyde (Merck, Darmstadt, Germany) in 0.1M cacodylate buffer, pH 7,4 subsequently post-fixed for 2h in 1% aqueous osmium tetroxide (Plano, Germany), dehydrated stepwise in a graded ethanol series and afterwards embedded in Epon 812 (Fluka, Buchs, Switzerland). Ultrathin (70-nm) sections were prepared with an ultramicrotome (EM UC6, Leica, Wetzlar, Germany), stained for 30 min with 1% uranyl acetate and 20 min in 3% lead citrate. Sections were examined at 50 kV in a Zeiss 109 transmission electron microscope (Zeiss, Oberkochen, Germany).

### Sample preparation for LC-MS/MS

For the proteome analysis we collected and processed a total of ~12,600 oocytes or embryos from May 2014 to October 2016. Specifically, we lysed, in triplicate, an average of ~600 oocytes/embryos per developmental stage: unfertilized oocytes, fertilized oocytes with pronuclei, and preimplantation embryos at the 2-, 4-, 8-cell, advanced morula and blastocyst stages. The samples were true biological replicates that were handled independently from start to end. Protein quantification was performed with our established spike-in SILAC-based labeling pipeline (Pfeiffer et al. 2011; Pfeiffer et al. 2015; Wang et al. 2016). Briefly, oocytes and embryos were deprived of the zona pellucida by pipetting in warm acidic Tyrode solution for 30-60 seconds and then rinsing in protein-free HCZB medium (BSA replaced through *polyvinylpyrrolidone* 40 kDa). Each sample lysate was then mixed with an equal amount of isotopically labeled (heavy) lysate from an embryo-derived F9 cell line capable of teratoma formation (Berstine et al. 1973), digested with trypsin, and subjected to MS analysis.

F9 EC cells were grown for several passages in RPMI 1640 medium (PAA, Cölbe, Germany), supplemented with 10% dialyzed fetal calf serum (Sigma, Deisenhofen, Germany), heavy amino acids ^13^C_6_ ^15^N_2_-L-Lysine (K8) and ^13^C_6_ ^15^N_4_-L-Arginine (R10; Silantes, Martinsried, Germany) as well as Glutamine and the antibiotics penicillin and streptomycin (Gibco, Darmstadt, Germany). The extent of labeling was 97.8%. The F9 EC cell line was originally isolated by Bernstine *et al.* (Berstine et al. 1973) as a subline of the teratocarcinoma OTT6050, established by implanting a 6 day-old embryo in the testis of a 129/J mouse. Thus, F9 EC cells have many characteristics of early mouse embryonal cells (Alonso et al. 1991) and are expected to provide a labeled counterpart for a large share of the proteins present in early embryos.

### LC-MS/MS analysis of SILAC mixtures

Subsequent to the tryptic digest, the peptide mixtures were offline fractionated by high pH reversed phase chromatography with fraction concatenation. The resulting peptide pools were analyzed by MS on a Q-Exactive mass spectrometer. The MS proteomics data have been deposited to the ProteomeXchange Consortium (http://proteomecentral.proteomexchange.org) via the PRIDE partner repository (Vizcaino et al. 2013) with the accession number PXD007082 and are summarized in Supplemental Table S5.

### Basic processing of raw LC-MS/MS data (MaxQuant, Perseus)

Raw data were processed by MaxQuant Software (v1.5.3.8, Martinsried, Bavaria, Germany) involving the built-in Andromeda search engine (Cox and Mann 2008; Cox et al. 2011). MS/MS spectra were searched against the mouse UniprotKB database (version from Dec. 2015) concatenated with reversed sequence versions of all entries and supplemented with common contaminants (see Supplemental Methods). Primary quantification was performed using the heavy F9 lysate mix as an internal standard, and ratios between corresponding light (L) and heavy (H) peptide versions were normalized to correct for unequal protein amounts and expressed as L/H (i.e., light/heavy: sample/SILAC internal standard). All these protein ratios are the means of at least two (light and heavy) peptide ratios from the raw spectra. Quality control determined that the sample corresponding to the blastocyst stage for replicate 3 was of low quality; this sample was therefore omitted from all analyses. The ID mapping procedure in some cases returned more than one gene name for a given peptide group; those may or may not correspond to distinct genes. To avoid ambiguities, we excluded such entries from the dataset.

### Protein data normalization and batch correction

We log2-transformed and quantile-normalized the L/H ratios of all proteins detected at least in two developmental stages in at least two replicates. To correct for the batch effect (see Supplemental Fig. S22), we performed an ANOVA for each protein, using the log_2_-transformed L/H ratios as response variable and the replicate identifier as categorical explanatory variable *X_i_*:

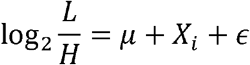

where *μ* is the global mean for the protein and denotes the error. The residuals of the model were used as the corrected L/H ratios for each protein, after adding to each value the global mean μ for the given protein as a constant. Batch-corrected, normalized L/H ratios were used to express protein abundance throughout this study.

### RNA isolation and RNA sequencing

For the transcriptome analysis we collected and lysed, in duplicate, an average of 214 oocytes/embryos per developmental stage: unfertilized oocytes, fertilized oocytes with pronuclei and preimplantation embryos at the (early and late) 2-, 4-, 8-cell, advanced morula and blastocyst stages, on which we then performed RNA sequencing (RNA-seq). Total RNA was converted to cDNA using the Smarter system (Takara) and sequencing libraries were prepared using the Nextera kit (Illumina). Libraries were sequenced on Illumina HiSeq 3000 platform to obtain ~43 million 36-base-single-end reads per library. The raw data are available at the DNA Databank of Japan (DDBJ) Sequence Read Archive (DRA005956 and DRA006335).

### RNA-seq trimming and mapping

Low quality reads were filtered using Trimmomatic (version 0.36, (Bolger et al. 2014)) with the following parameters: HEADCROP:15 LEADING:3 TRAILING:3 SLIDINGWINDOW:4:15 MINLEN:20. The remaining reads were mapped to the *Mus musculus* Ensembl GRCm38 assembly using TopHat (version 2.1.1, (Kim et al. 2013)) and Bowtie (version 2.2.9, (Langmead and Salzberg 2012)). As the only non-default parameter for TopHat, we provided the GRCm38 Ensembl 87 (version 1) GTF annotation with the “-G” option. The number of reads mapped to each gene was quantified with with HTSeqCount (version 0.6.1, (Anders et al. 2015)) using standard parameters.

### RNA differential expression analysis

A matrix containing the number of reads mapped to each protein-coding gene for each sample was used as input for differential expression analysis with the DESeq2 R/Bioconductor package (Anders and Huber 2010; Love et al. 2014). The P-values obtained from DESeq2 were adjusted with Benjamini-Hochberg’s method to control the false discovery rate (FDR) (Benjamini and Hochberg 1995). Genes were considered significantly differentially expressed on the basis of (log_2_) fold-change ((log_2_) fold-change ≥1 or ≤−1 between the two developmental stages considered) and FDR≤1×10^-5^. Expression values of protein-coding transcripts were calculated using DESeq2 using the regularized log-transformation (Anders and Huber 2010; Love et al. 2014).

### Protein differential expression analysis

For each protein detected at least in two developmental stages in at least two replicates we computed a linear model:

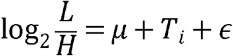

where *μ* is the global mean for the gene, *T_i_* is a categorical explanatory variable representing the developmental stage, and *ϵ* denotes the error. For 1,290 proteins, the ANOVA P-value corresponding to *T_i_* was 0.05.

### Validation of proteins by enzymatic assays and immunofluorescence

Results of enzymatic assays for G6PD (EC 1.1.1.49) and HPRT (EC 2.4.2.7) were retrieved from the literature (Brinster 1966; Epstein et al. 1969; Epstein 1970; Kratzer and Gartler 1978; Ayabe et al. 1994).

Additional proteins including proteins without enzymatic activity were verified by immunofluorescence, using commercial antibodies. For each target gene, at least 5 MII oocytes or embryos per stage were examined using the following antibodies, all rabbit polyclonal: anti-DNAJB11 (Sigma-Aldrich cat.no. HPA010814), anti-PDIA3 (Abcam cat.no. ab228789), anti-TOP1 (Sigma-Aldrich cat.no. HPA019039), anti-Rc3h1 (Thermo Scientific catalog no. PA5-34519), anti-Alppl2 (Thermo Scientific catalog no. PA5-22336), anti-DDX6 (Thermo scientific catalog no. PA5-55012). Secondary antibodies were Alexa-Fluor conjugates reactive against the species of the primary antibody. Following our standard fixation, permeabilization, incubation and washing protocol (Schwarzer et al. 2012), samples were imaged using a 20X objective on an inverted motorized Nikon TiE2000 microscope fitted with an Andor Dragonfly spinning disc confocal unit Scanning System. Immunofluorescent signals were quantified using Image-J (Schneider et al. 2012).

For each protein, we calculated the Spearman’s rank correlation coefficient between the immunofluorescent signals or enzymatic measurements and the average L/H ratios in our dataset for all available developmental stages. For proteins for which multiple sets of measurements were available we computed and considered as many correlation coefficients. An empirical P-value was computed by randomly associating each of the protein measurements from the literature with one of the corresponding sets of measurements in our dataset (see Supplemental Table S3) and repeating this 10,000 times. The reported empirical P-value is the number of times in which we obtained the same number of correlation coefficients greater or equal than 0.6 as with the original data out of the 10,000 attempts, expressed as a relative frequency.

### TaqMan validation of RNAseq

For each target gene, the cDNA equivalent of 10 MII oocytes or embryos per stage was used. Total RNA was isolated from large pools (>100 oocytes or embryos) using Quick-RNA™ MicroPrep (Zymo Research) following the manufacturer’s instructions and was reverse-transcribed on a GeneAmp^®^ PCR System 9700 (Applied Biosystems). Real-time quantitative PCR reactions were performed on cDNA on a 7900 HT FAST Realtime PCR System (Applied Biosystems). PrimeTime^®^Predesigned qPCR Assay (6-FAM/ZEN/IBFQ) from Integrated DNA Technologies were used. Assay IDs: Dnajb11_Mm.PT.58.9272431, Pdia3_Mm.PT.8194853; Top1_Mm.PT.58.6752545. All samples were processed as technical duplicates/replicates. Data were analyzed using the Applied Biosystems RQ Manager (Version 1.2.2) and Microsoft Excel.

### Data Access

The proteomic data from this study have been submitted to the ProteomeXchange Consortium (http://proteomecentral.proteomexchange.org) via the PRIDE partner repository (Vizcaino et al. 2013) under accession number PXD007082. The sequence data generated for this study have been submitted to DNA Databank of Japan (DDBJ, http://www.ddbj.nig.ac.jp/) under the accession numbers DRA005956 and DRA006335.

## Supporting information

## Acknowledgements

We thank the Max-Planck-Institute for Molecular Biomedicine and its Director, Prof. Hans R. Schöler, for infrastructural support. We thank the personnel of the MPI mouse housing facility for making it possible to collect as many oocytes and embryos as needed for the proteomics and RNA sequencing. We are indebted to Annalen Nolte for processing the probes for mass spectrometry, and Terumi Horiuchi for processing the probes for RNA sequencing. Jeroen Krijgsveld commented on an earlier version of the manuscript. This study was supported by the Deutsche Forschungsgemeinschaft (grant DFG BO 2540/4-3 to M.B., grant TA 1076/1-1 to L.T., and grant FU-583/5-1 to G.F.; O.E.P. acknowledges support for the transmission electron microscopy from the SFB 944).

## Author contributions

L.T., G.F. and M.B. planned the study. H.C.D. performed the proteomics experiments. Y.S. performed the RNA-seq experiments. S.I., M.E., L.T., G.F. and M.B. designed the analytical pipeline, analyzed and interpreted the data. L.T., G.F. and M.B. wrote the manuscript with help from S.I. and M.E. O.E.P. performed the transmission electron microscopy. S.I. and E.C. performed the confocal immunofluorescence imaging. W.M. provided intellectual guidance with RNA-seq analysis and feedback on the experimental design. All authors discussed the results and commented on the manuscript.

