## Supplementary material for "An integrated genome-wide multi-omics analysis of gene expression dynamics in the preimplantation mouse embryo"

**Running Title:** Multi-omics of the preimplantation mouse embryo

**Keywords:** Preimplantation development, Proteome, Transcriptome, Model Organism

### Supplementary Figures

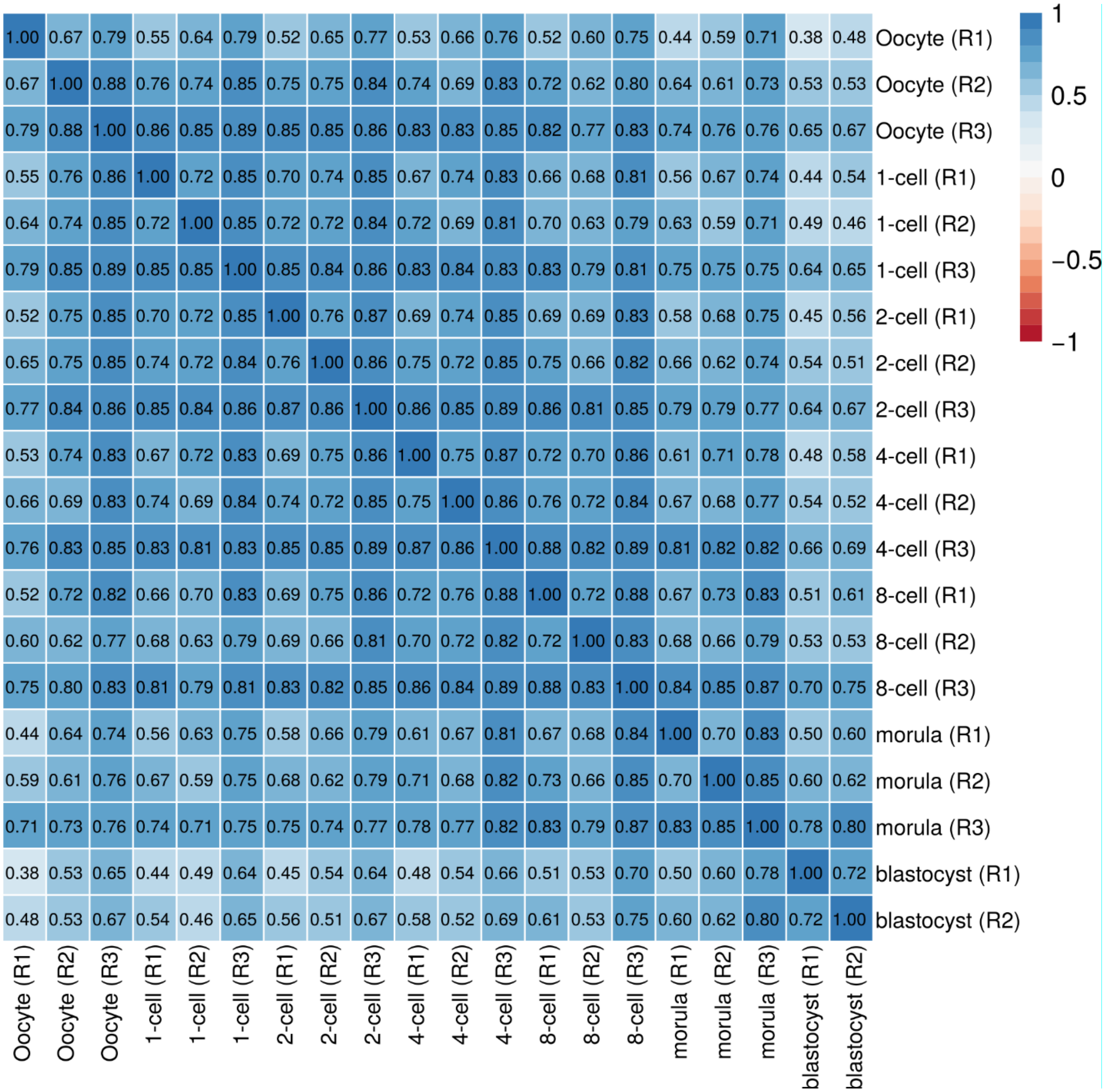

**Supplemental Fig. S1. Proteome replicates are strongly correlated.** Spearman's rank correlations of pairwise comparisons between the protein ($log_2$) L/H ratios of the 20 samples in the study.

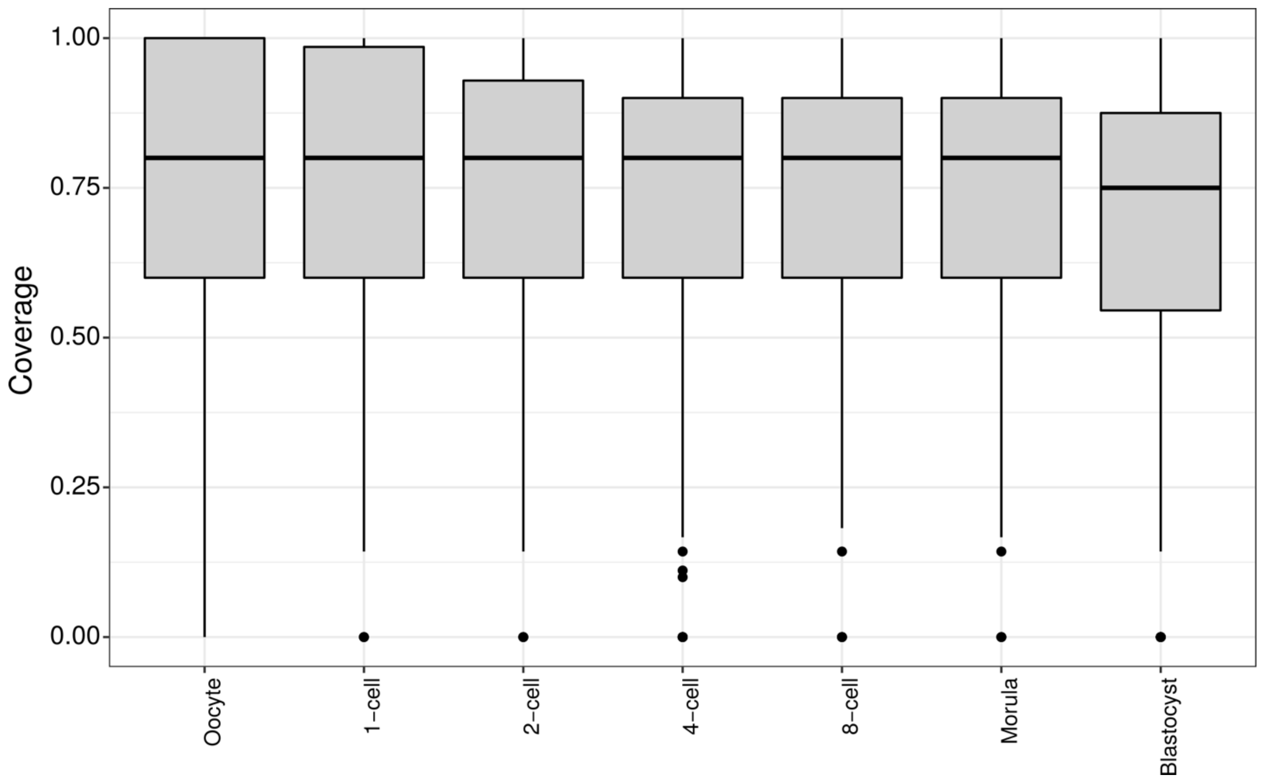

**Supplemental Fig. S2. Coverage distribution for 233 protein complexes across different development stages.** Complexes comprise 5 to 153 protein members (with a median of 7). The coverage was computed as the fraction of the members of a complex detected in at least one of the replicates of the developmental stage under consideration. The distributions of the fractions are shown as boxplots.

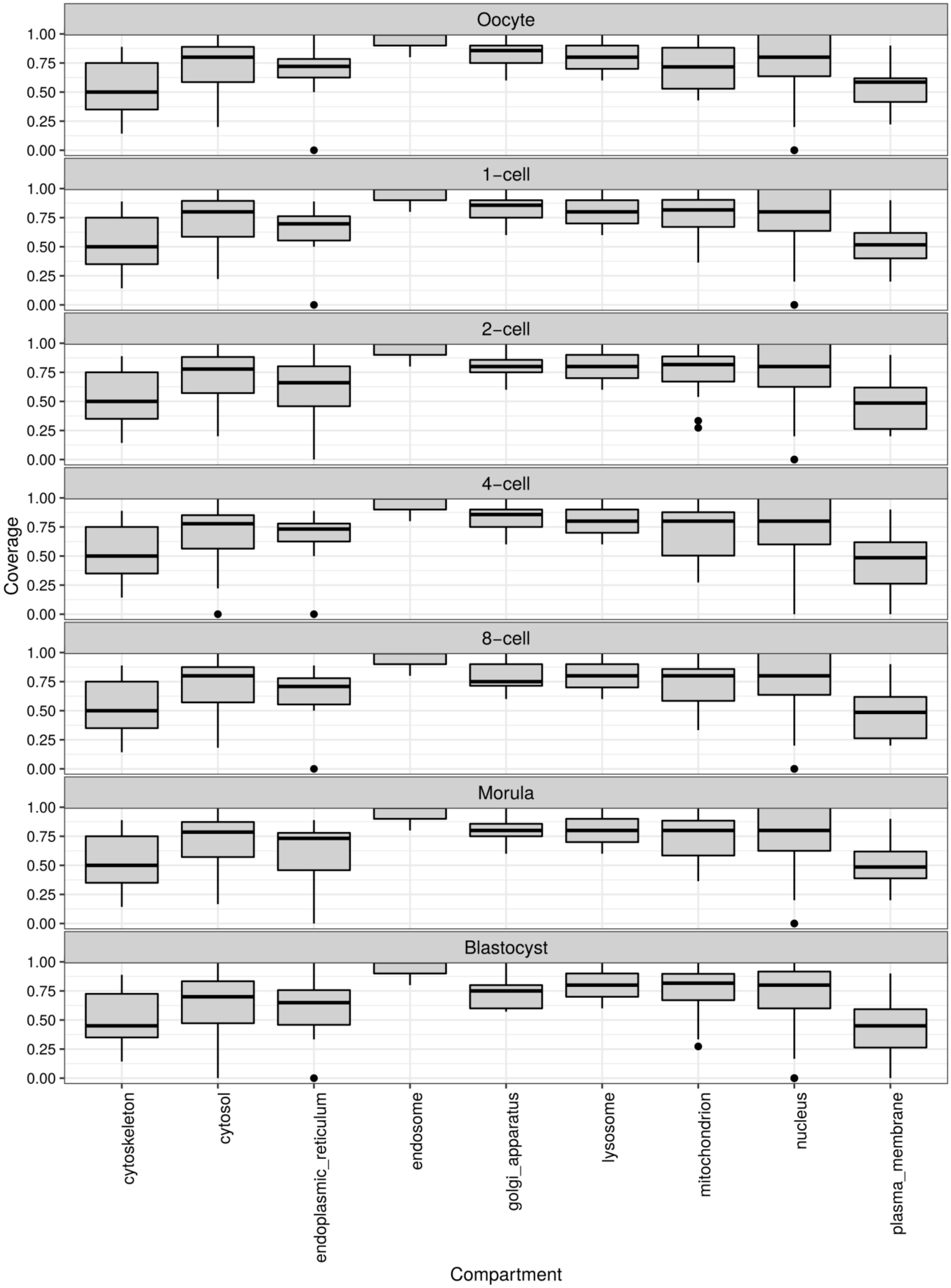
Oocyte
1-cell
2-cell
4-cell
8-cell
Morula
Blastocyst
Coverage
1.00
0.75
0.50
0.25
0.00
cytoskeleton
cytosol
endoplasmic_reticulum
endosome
golgi_apparatus
lysosome
mitochondrion
nucleus
plasma_membrane
Compartment

**Supplemental Fig. S3. Coverage distribution for 233 protein complexes, according to their subcellular localization, across different development stages (pooled replicates).** Complexes comprise 5 to 153 protein members (with a median of 7). Each complex was assigned the subcellular localization(s) of the largest group(s) of proteins among its members, as annotated in the Compartments database (https://compartments.jensenlab.org/, (Binder et al. 2014)). Thus, each subcellular localization was associated with 2 to 153 complexes (with a median of 8). The coverage was computed as the fraction of the members of a complex detected in at least one of the replicates of the developmental stage under consideration. The distributions of the fractions are shown as boxplots.

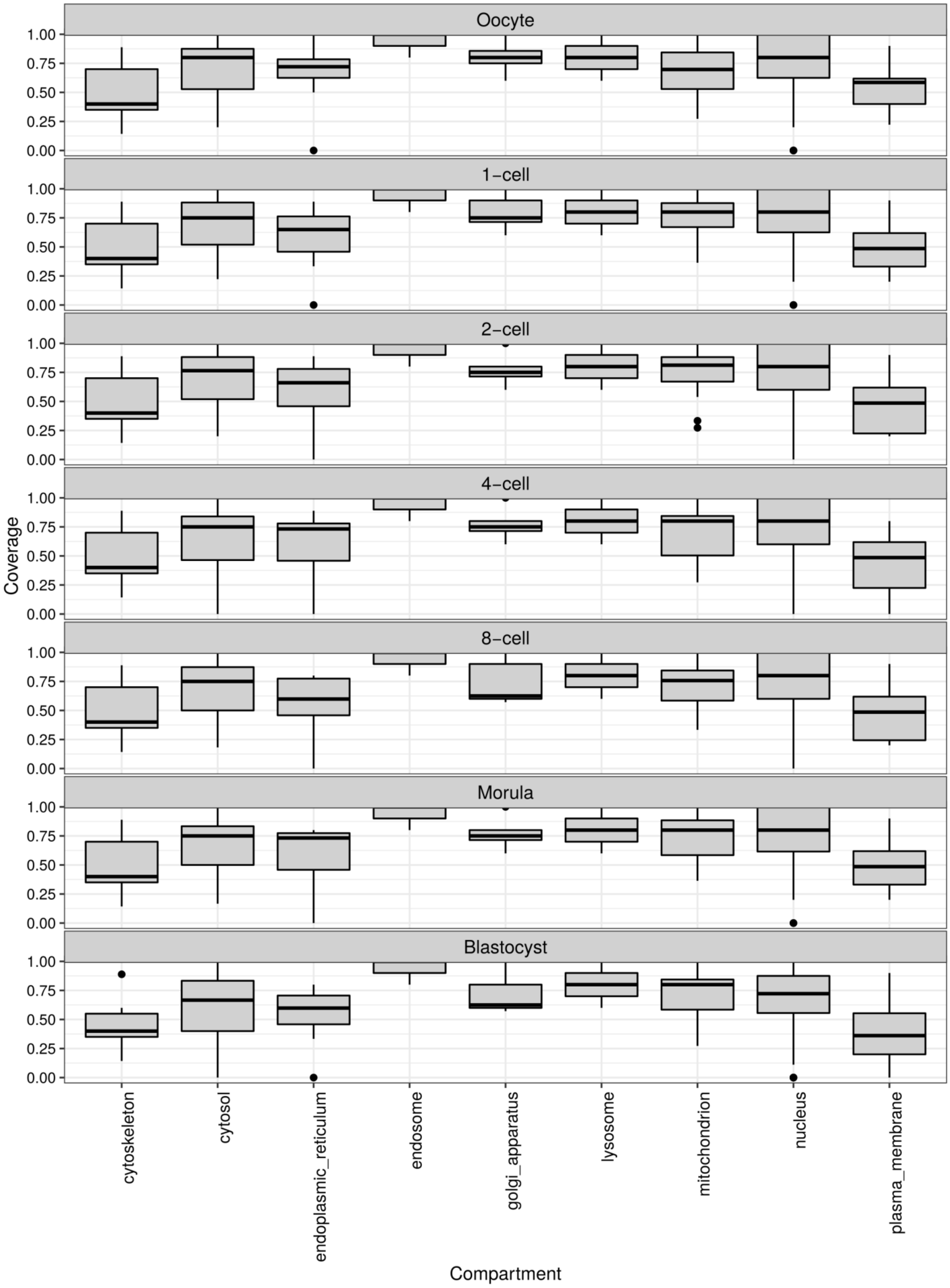

Oocyte
1-cell
2-cell
4-cell
8-cell
Morula
Blastocyst
1.00
0.75
0.50
0.25
0.00
Coverage
cytoskeleton
cytosol
endoplasmic_reticulum
endosome
golgi_apparatus
lysosome
mitochondrion
nucleus
plasma_membrane
Compartment

**Supplemental Fig. S4. Coverage distribution for 233 protein complexes, according to their subcellular localization, across different development stages (replicate 1).** The coverage was computed as the fraction of the members of a complex detected in replicate “1” of the developmental stage under consideration. See Supplemental Fig. S3 for details.

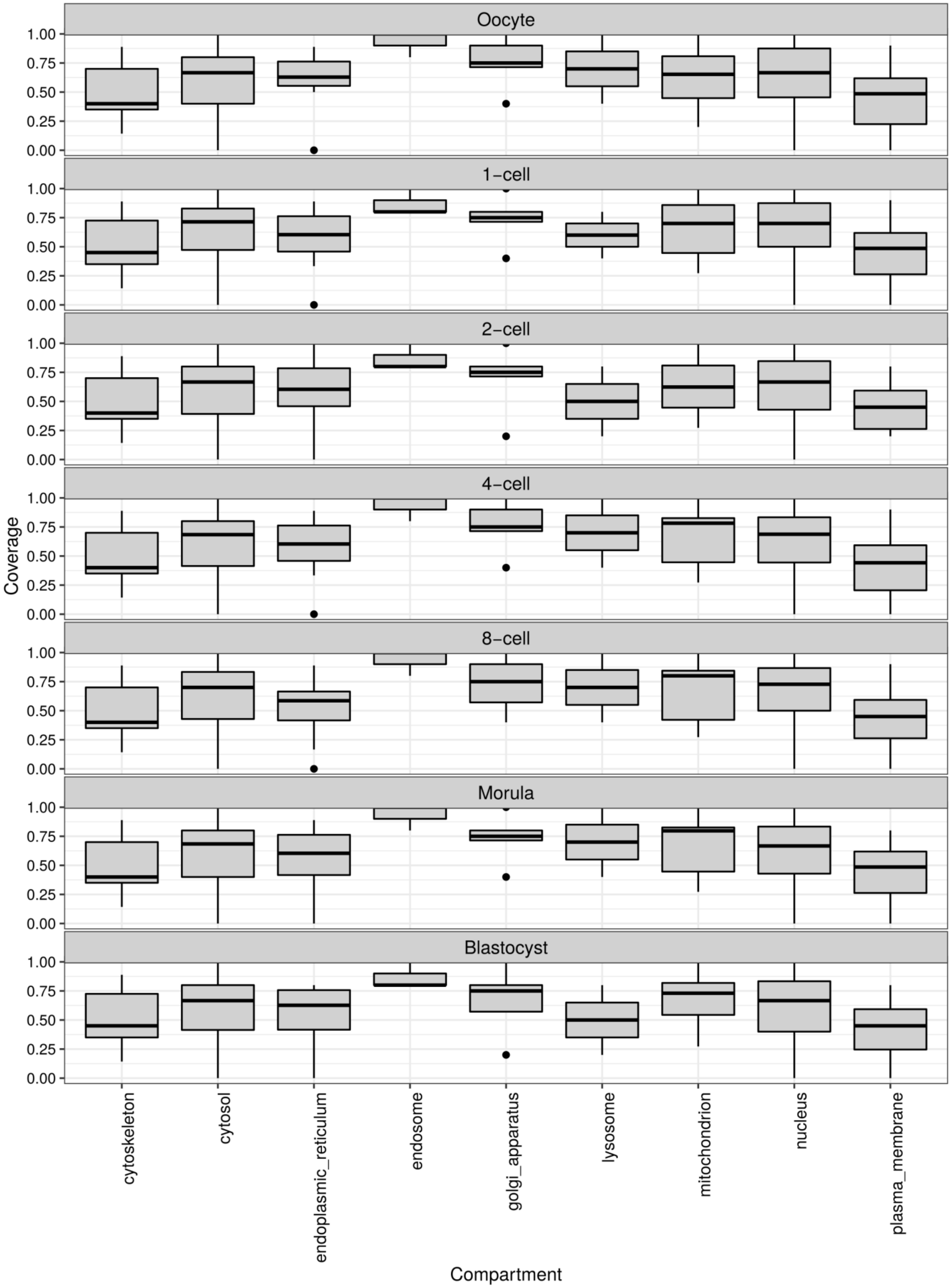
Oocyte
1-cell
2-cell
4-cell
8-cell
Morula
Blastocyst
Coverage
0.00
0.25
0.50
0.75
1.00
cytoskeleton
cytosol
endoplasmic_reticulum
endosome
golgi_apparatus
lysosome
mitochondrion
nucleus
plasma_membrane
Compartment

**Supplemental Fig. S5. Coverage distribution for 233 protein complexes, according to their subcellular localization, across different development stages (replicate 2).** The coverage was computed as the fraction of the members of a complex detected in replicate “2” of the developmental stage under consideration. See Supplemental Fig. S3 for details.

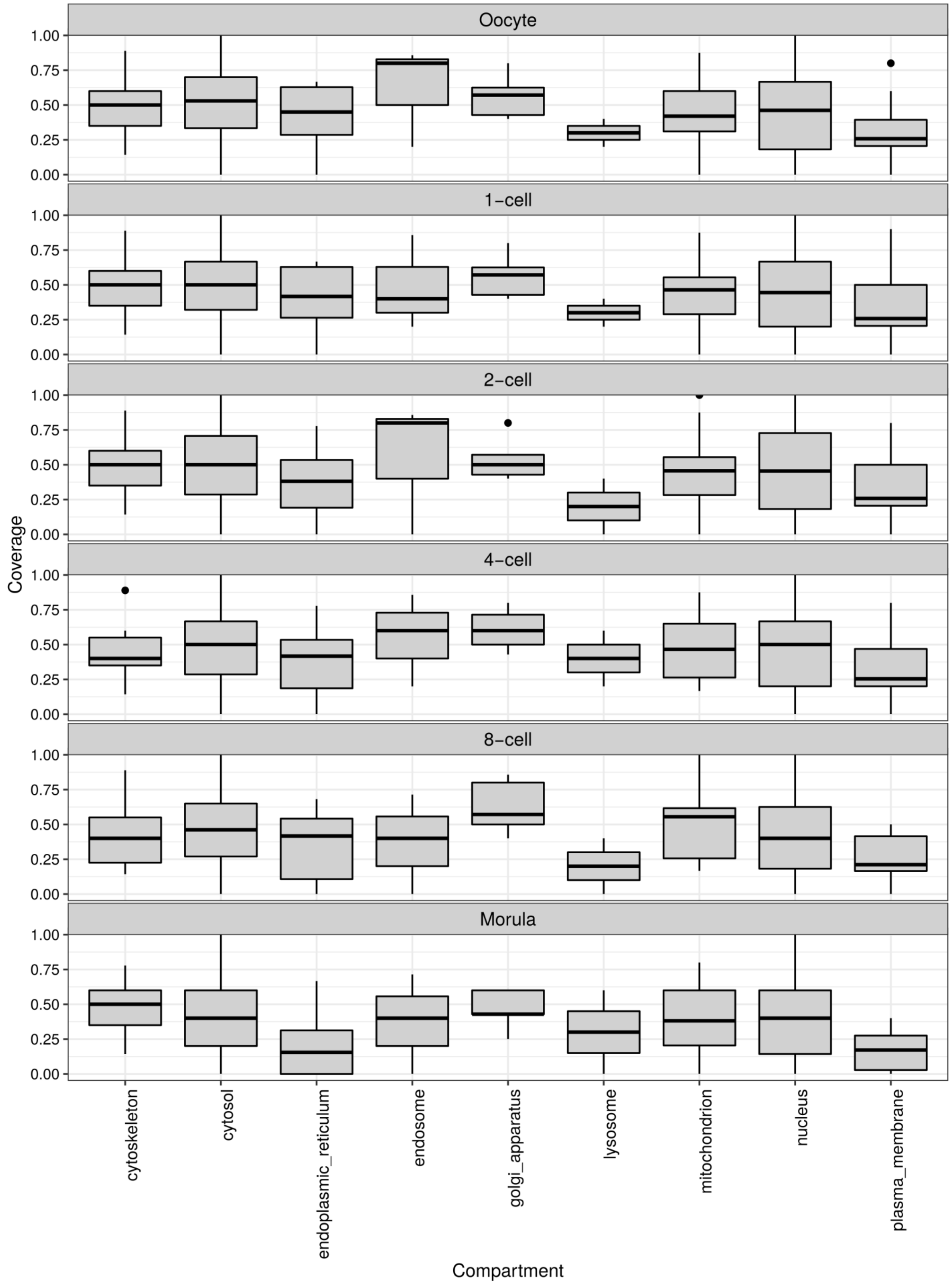

Oocyte
1-cell
2-cell
4-cell
8-cell
Morula
Coverage
1.00
0.75
0.50
0.25
0.00
cytoskeleton
cytosol
endoplasmic_reticulum
endosome
golgi_apparatus
lysosome
mitochondrion
nucleus
plasma_membrane
Compartment

**Supplemental Fig. S6. Coverage distribution for 233 protein complexes, according to their subcellular localization, across different development stages (replicate 3).** The coverage was computed as the fraction of the members of a complex detected in replicate “3” of the developmental stage under consideration. See Supplemental Fig. S3 for details.

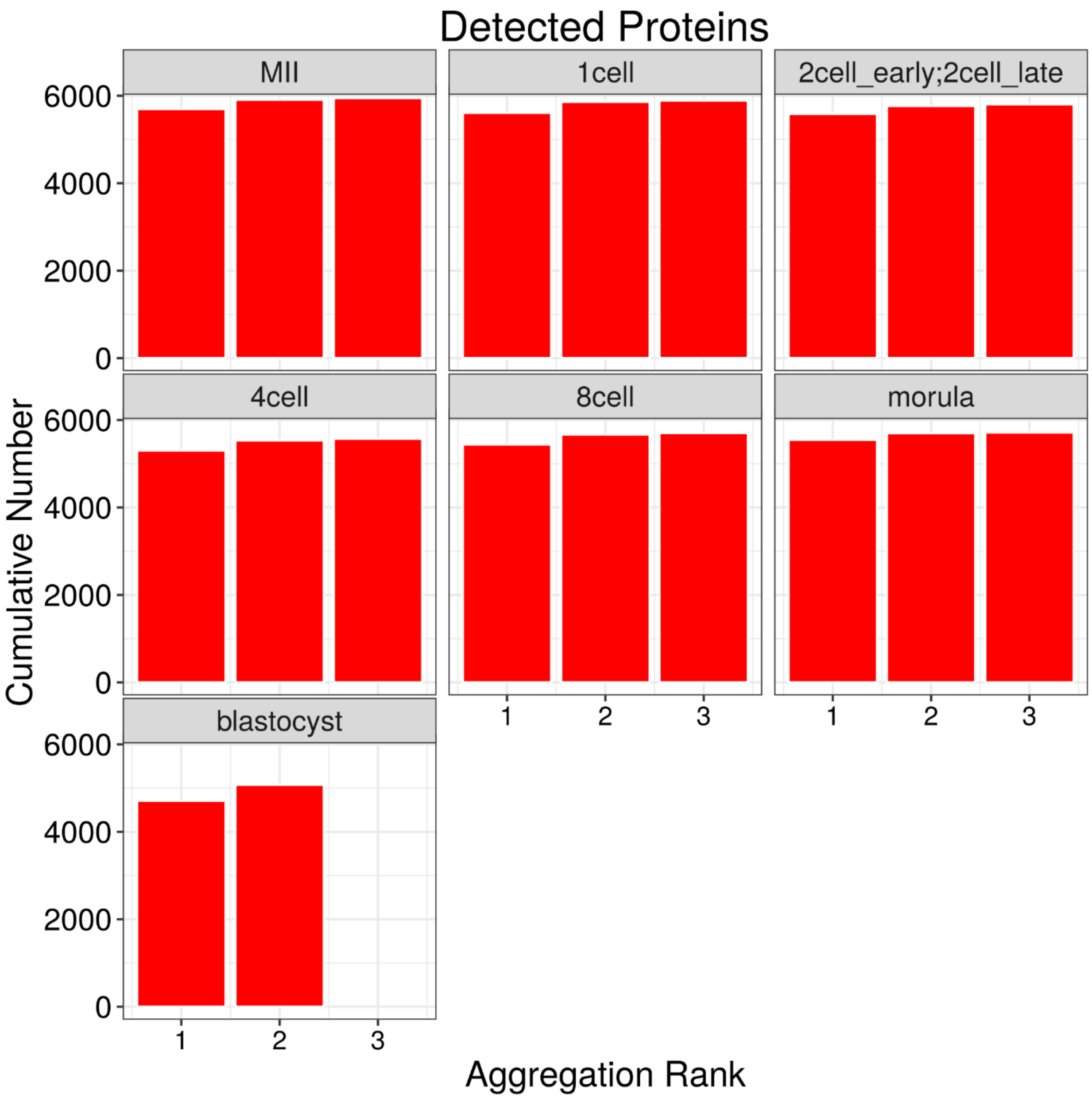

**Supplemental Fig. S7. Cumulative number of detected proteins, with an arbitrary ordering of the replicates.** Proteome coverage rapidly saturates with the aggregation of replicates at ~5,500 to ~6,000, depending on the developmental stage. This indicates that proteome coverage is likely not to increase much more by adding replicates quantified using SILAC.

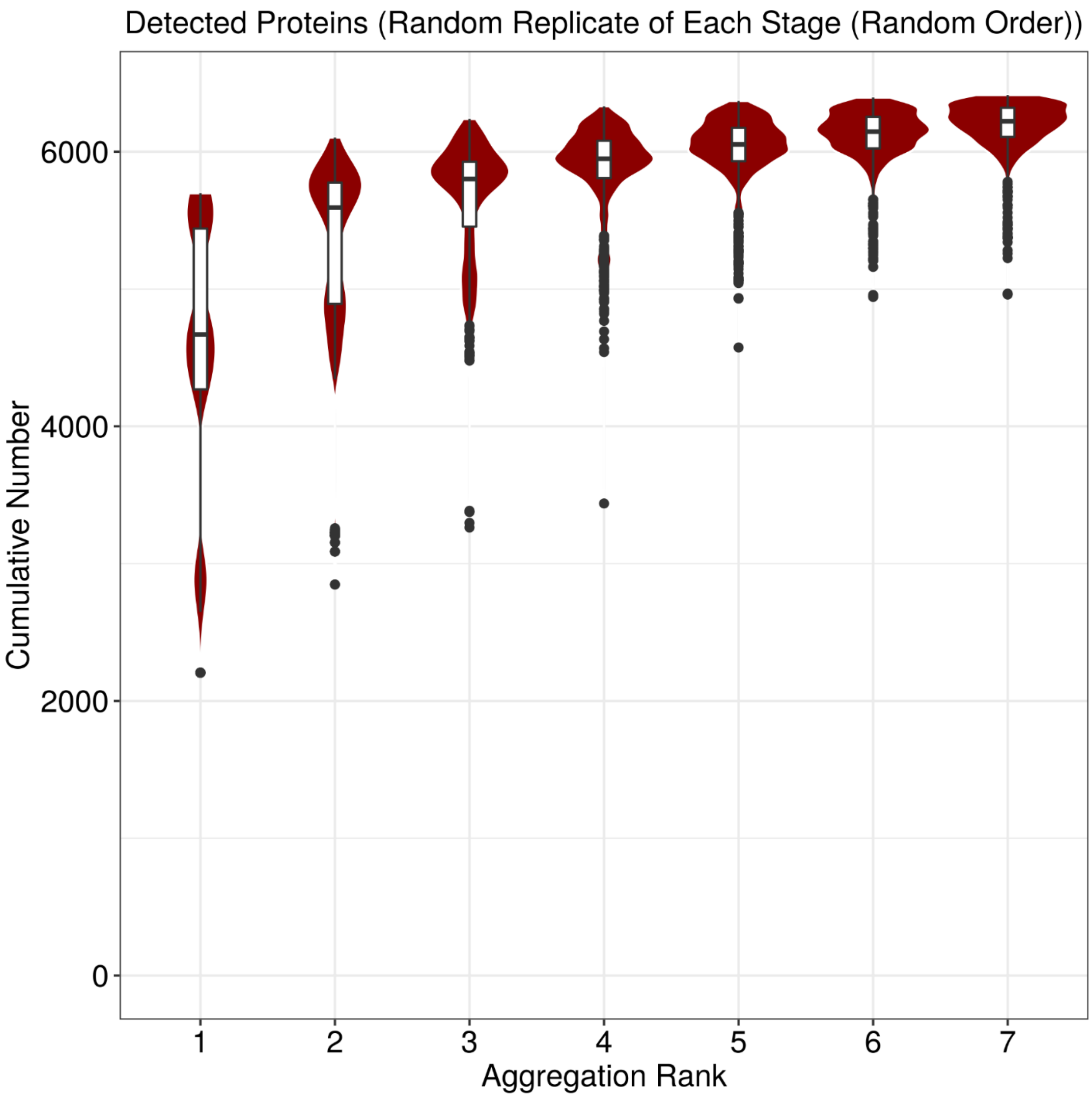

**Supplemental Fig. S8. Cumulative number of detected proteins, with an arbitrary ordering of the developmental stages.** The data summarizes 1,000 series constructed by sequentially selecting one random replicate from a random developmental stage, so that each developmental stage is represented exactly once. The violin plot at each aggregation rank represents the distribution of the cumulative number of detected proteins, computed for each series. Proteome coverage saturates rapidly, such that the aggregation of the last four samples results in a very small increase of the median cumulative number of detected proteins. This indicates that proteome coverage is likely not to increase much more by adding similar samples quantified using SILAC.

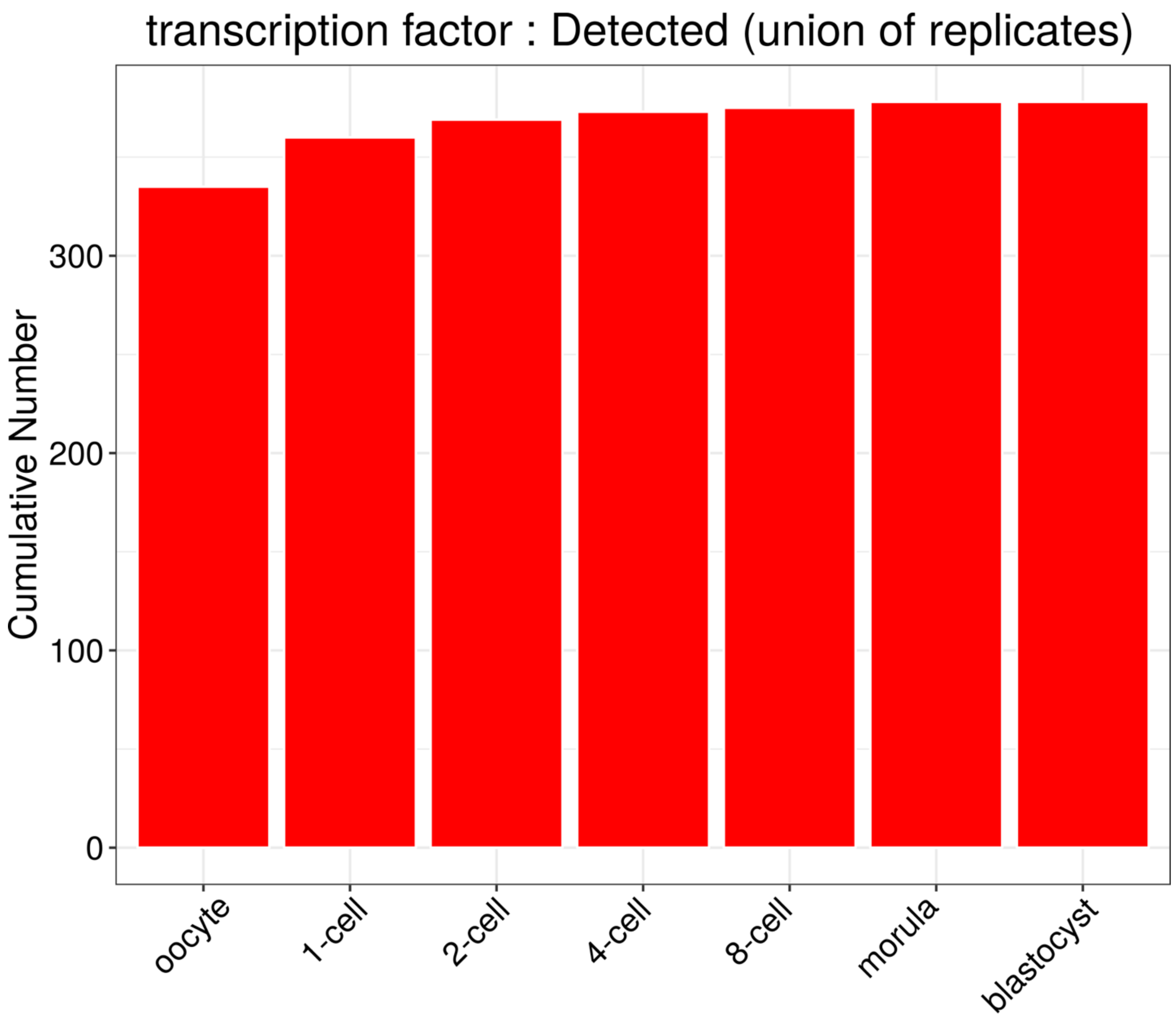

**Supplemental Fig. S9. Cumulative number of transcription factor (TF) proteins detected in the union of the replicates as preimplantation development progresses**. TFs were identified based on the PANTHER Classification System (http://www.pantherdb.org/, (Mi et al. 2013; Mi et al. 2017)) as all proteins annotated with the class PC00218 ("transcription factor") and its children and descendants in the PANTHER hierarchy. 1,392 proteins among the complete mouse proteome can be classified as "transcription factors".

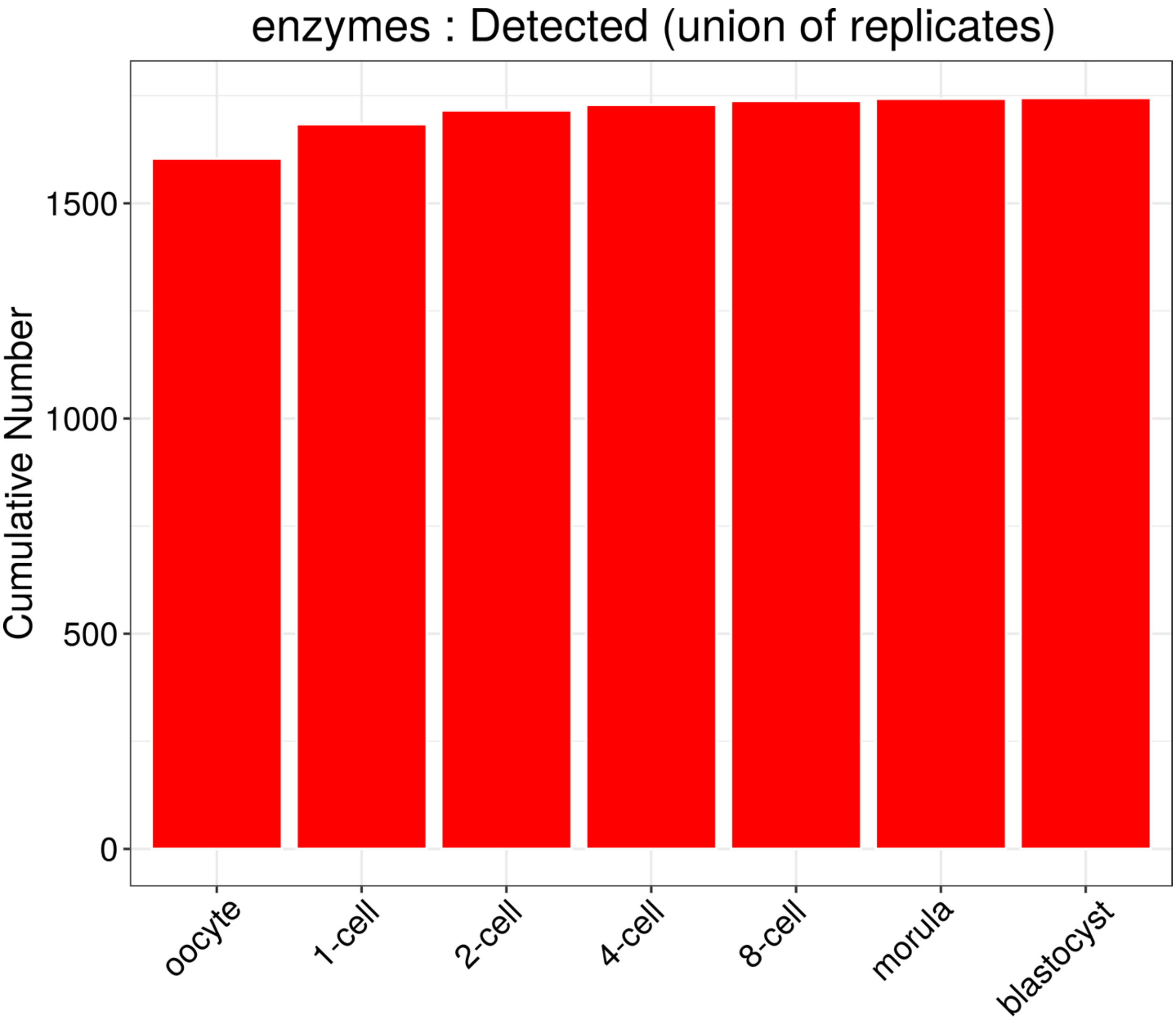

**Supplemental Fig. S10. Cumulative number of enzymes detected in the union of the replicates as preimplantation development progresses**. Enzymes were defined as proteins comprised in PANTHER classes (http://www.pantherdb.org/, (Mi et al. 2013; Mi et al. 2017)) with names including the "-ase" suffix and its children and descendants. 3,988 proteins among the complete mouse proteome can be classified as "enzymes" in this manner.

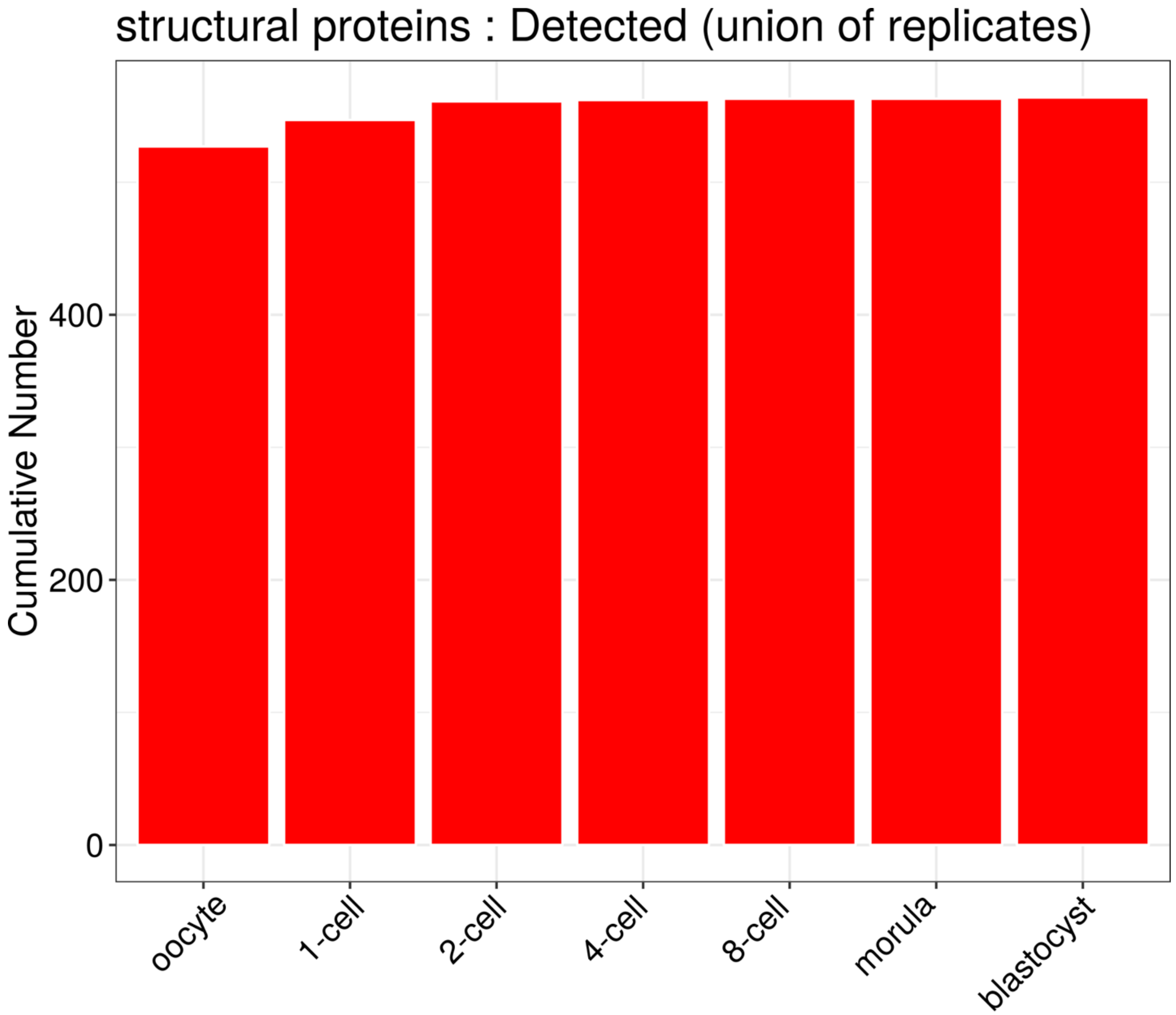

**Supplemental Fig. S11. Cumulative number of structural proteins detected in the union of the replicates as preimplantation development progresses.** Structural proteins were defined as proteins comprised in the PANTHER (http://www.pantherdb.org/, (Mi et al. 2013; Mi et al. 2017)) classes "cytoskeletal protein" (PC00085), "cell junction protein" (PC00070), "structural protein" (PC00211), "membrane traffic protein" (PC00150), "extracellular matrix protein" (PC00102), "cell adhesion molecule" (PC00069) and "viral coat protein" (PC00236) and all PANTHER classes that are children and descendants of these classes in the hierarchy of the PANTHER database. 1,875 proteins among the complete mouse proteome can be classified as “structural proteins” in this manner. See Supplemental Fig. S8 for details.

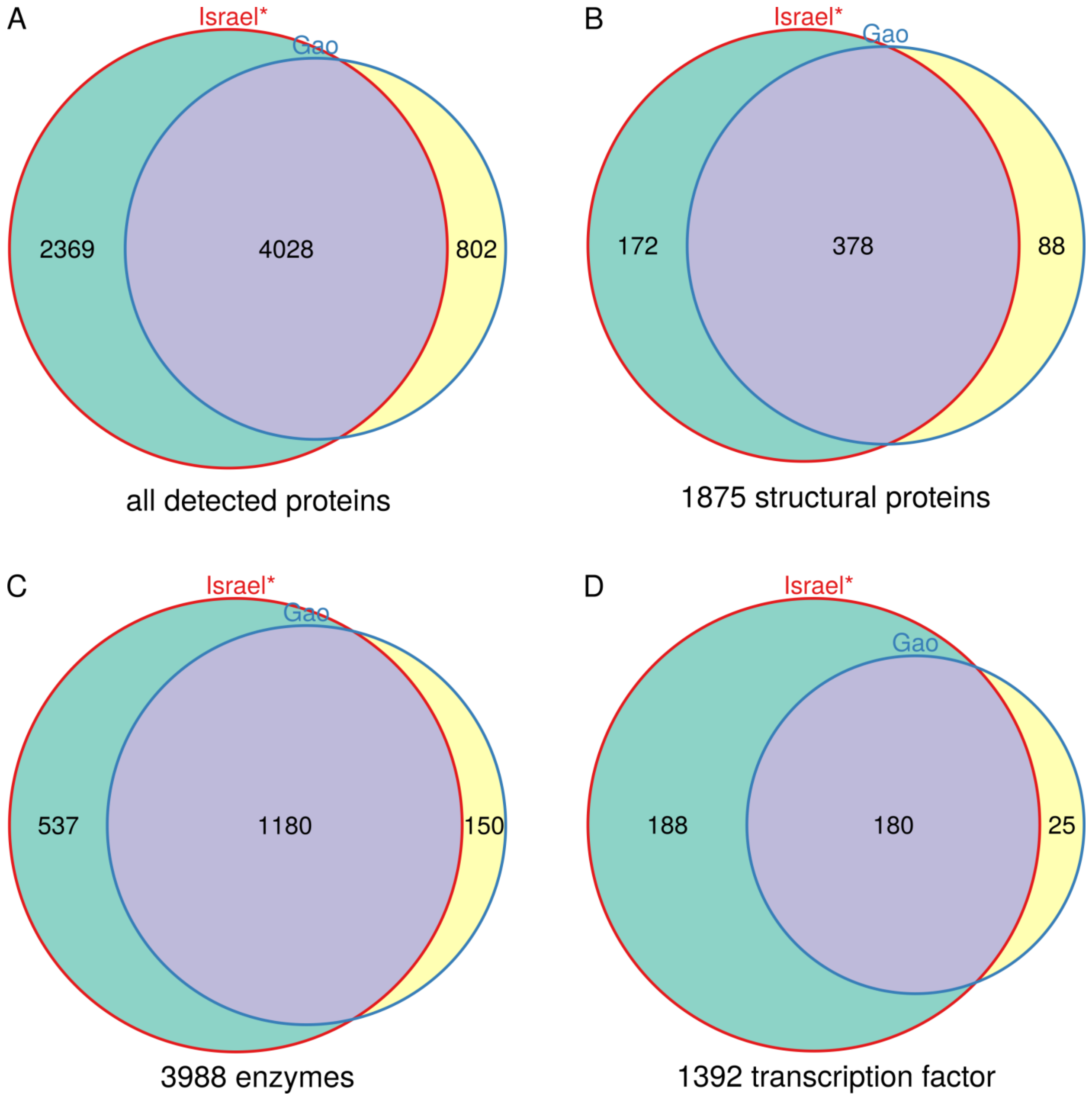

**Supplemental Fig. S12. Proteins detected in at least one replicate of at least one developmental stage (zygote to blastocyst only) in our dataset and in Gao et al's dataset (Gao et al. 2017)**. A) all proteins. B-C) Structural proteins, enzymes and transcription factors, defined based on the annotation of the PANTHER Classification System (http://www.pantherdb.org/, (Mi et al. 2013; Mi et al. 2017)). Structural proteins (B) were defined as proteins comprised in the PANTHER classes "cytoskeletal protein" (PC00085), "cell junction protein" (PC00070), "structural protein" (PC00211), "membrane traffic protein" (PC00150), "extracellular matrix protein" (PC00102), "cell adhesion molecule" (PC00069) and "viral coat protein" (PC00236) and all PANTHER classes that are children and descendants of these classes in the hierarchy of the PANTHER database. 1,875 proteins among the complete mouse proteome can be classified as "structural proteins" in this manner. Enzymes (C) were defined as proteins comprised in PANTHER classes with names including the "-ase" suffix and its children and descendants. 3,988 proteins among the complete mouse proteome can be classified as "enzymes" in this manner. Transcription factors (D) were defined as proteins comprised in the PANTHER class "transcription factor" (PC00218) and all its children and descendants. 1,392 proteins among the complete mouse proteome can be classified as "transcription factors" in this manner. The Venn diagram (generated using version 3.0 of the Vennerable R package) indicates the number of proteins detected in our dataset (Israel*) and in Gao et al's dataset (Gao)(Gao et al. 2017). Note that we excluded our oocyte data for comparability.

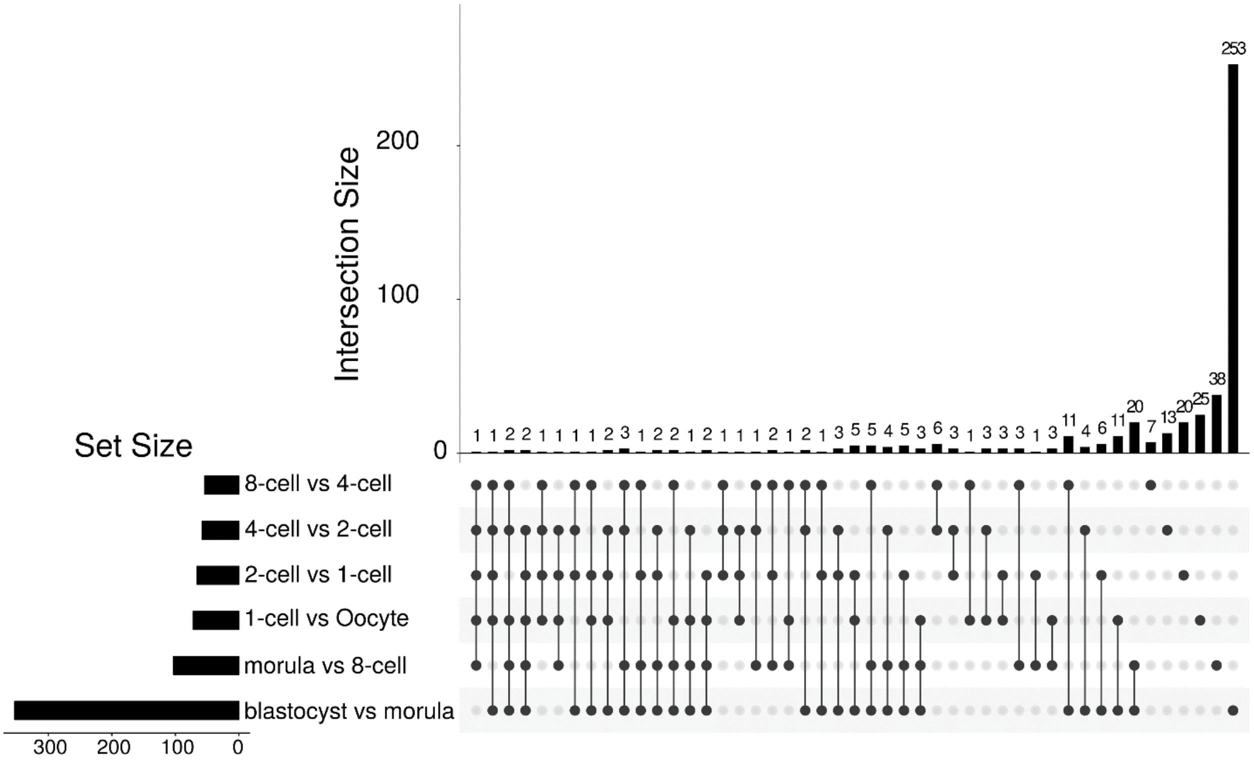

**Supplemental Fig. S13. UpSet plots (Lex et al. 2014) showing the overlap between the sets of differentially expressed proteins between pairs of consecutive developmental stages**. The bar represents the number of proteins shared by the pairs of stages indicated by the black dots and not by the stages indicated by the gray dots. A total of 488 proteins was differentially expressed between pairs of consecutive developmental stages; 356 were exclusively differentially expressed in one transition.

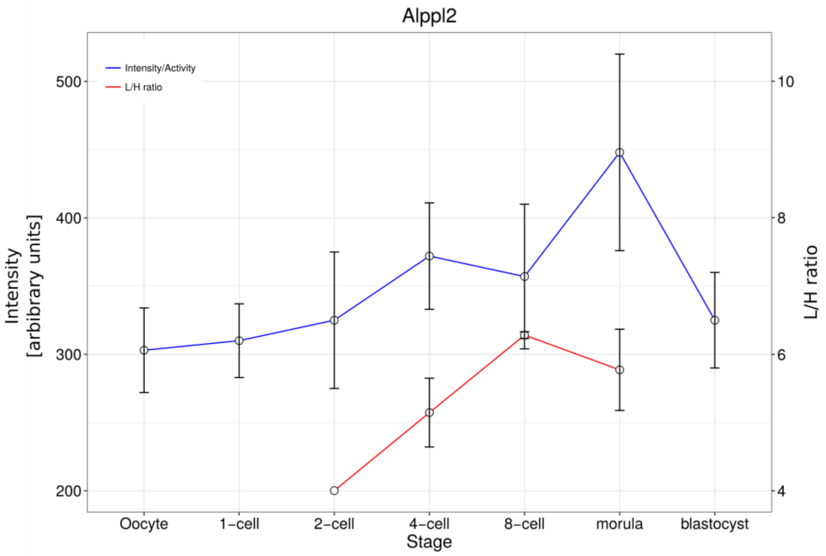
Alppl2
Intensity/Activity
L/H ratio
Intensity [arbibrary units]
L/H ratio
500
400
300
200
10
8
6
4
Oocyte
1−cell
2−cell
4−cell
8−cell
morula
blastocyst
Stage

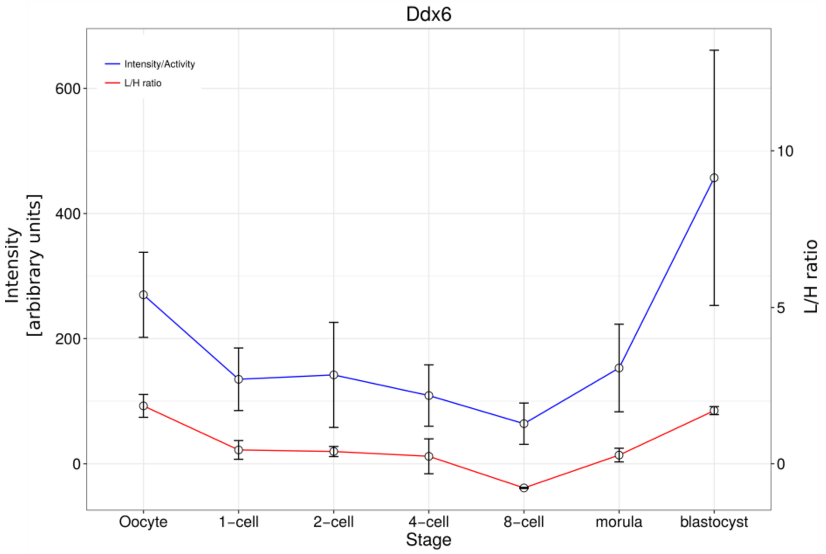
Ddx6
Intensity/Activity
L/H ratio
Intensity [arbibrary units]
L/H ratio
600
400
200
0
10
5
0
Oocyte
1−cell
2−cell
4−cell
8−cell
morula
blastocyst
Stage

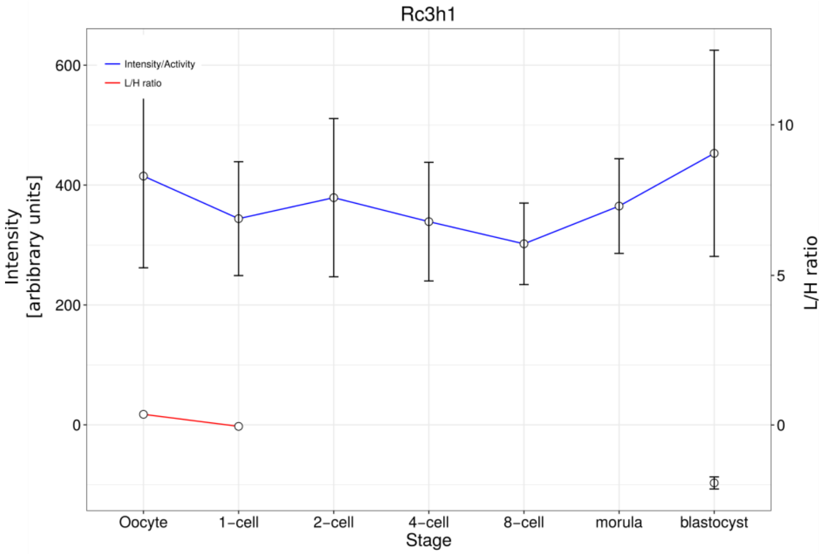
Rc3h1
Intensity/Activity
L/H ratio
Intensity [arbibrary units]
L/H ratio
600
400
200
0
10
5
0
Oocyte
1−cell
2−cell
4−cell
8−cell
morula
blastocyst
Stage

**Supplemental Fig. S14. Immunofluorescence validation of proteins levels.** Intensities of immunofluorescence images are in arbitrary units.

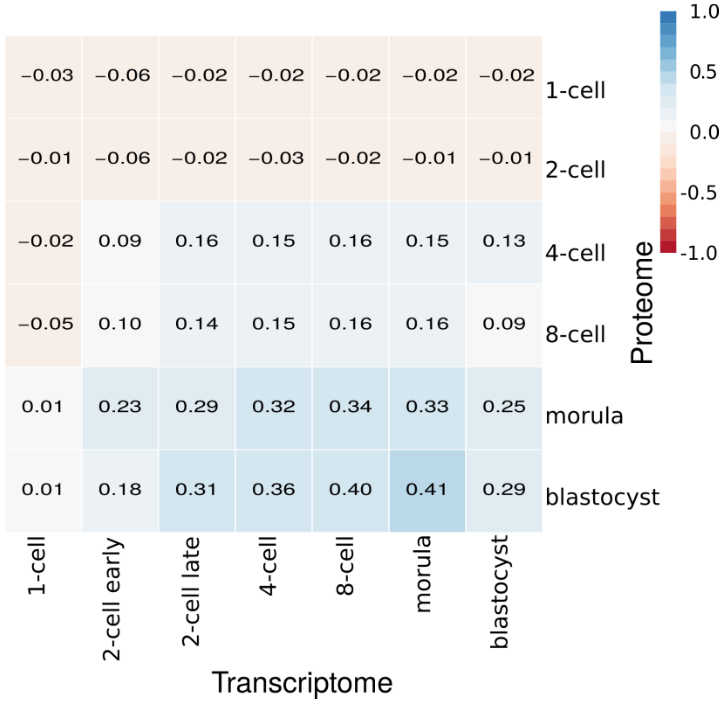

**Supplemental Fig. S15. Spearman's rank correlations between the fold-changes in $\log_2$ L/H ratios and expression values observed for the proteins and for their cognate transcripts, respectively, relative to the oocyte.** Sample group averages were considered for the calculation of pairwise correlations between the seven developmental stages in the study. Reported values are based on proteins detected in at least two replicates of each of the two developmental stages involved in each correlation calculation and their cognate transcript.

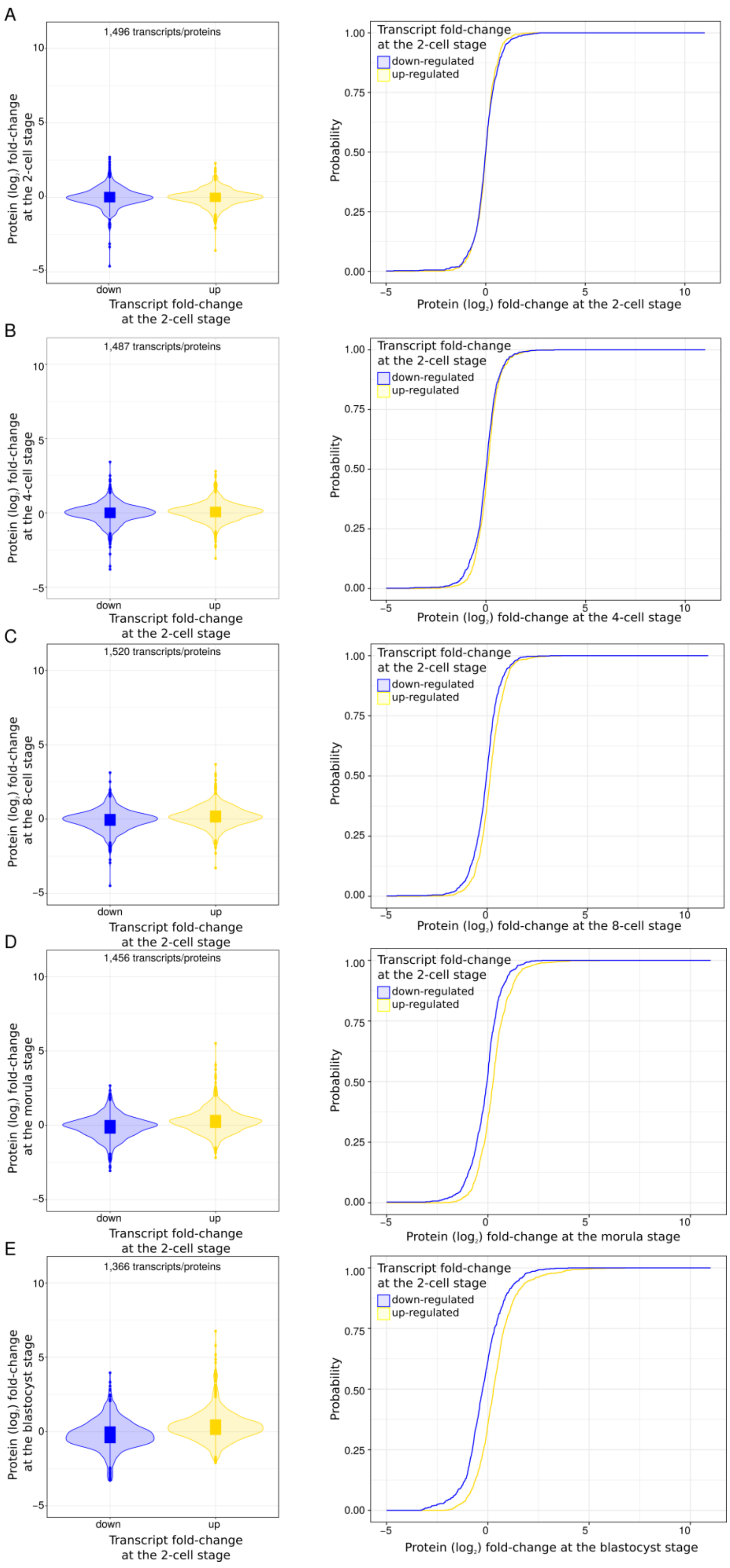
A
1,496 transcripts/proteins
Protein (log2) fold-change at the 2-cell stage
Transcript fold-change at the 2-cell stage
down
up
Transcript fold-change at the 2-cell stage
down-regulated
up-regulated
Probability
Protein (log2) fold-change at the 2-cell stage
B
1,487 transcripts/proteins
Protein (log2) fold-change at the 4-cell stage
Transcript fold-change at the 2-cell stage
Protein (log2) fold-change at the 4-cell stage
C
1,520 transcripts/proteins
Protein (log2) fold-change at the 8-cell stage
Transcript fold-change at the 2-cell stage
Protein (log2) fold-change at the 8-cell stage
D
1,456 transcripts/proteins
Protein (log2) fold-change at the morula stage
Transcript fold-change at the 2-cell stage
Protein (log2) fold-change at the morula stage
E
1,366 transcripts/proteins
Protein (log2) fold-change at the blastocyst stage
Transcript fold-change at the 2-cell stage
Protein (log2) fold-change at the blastocyst stage

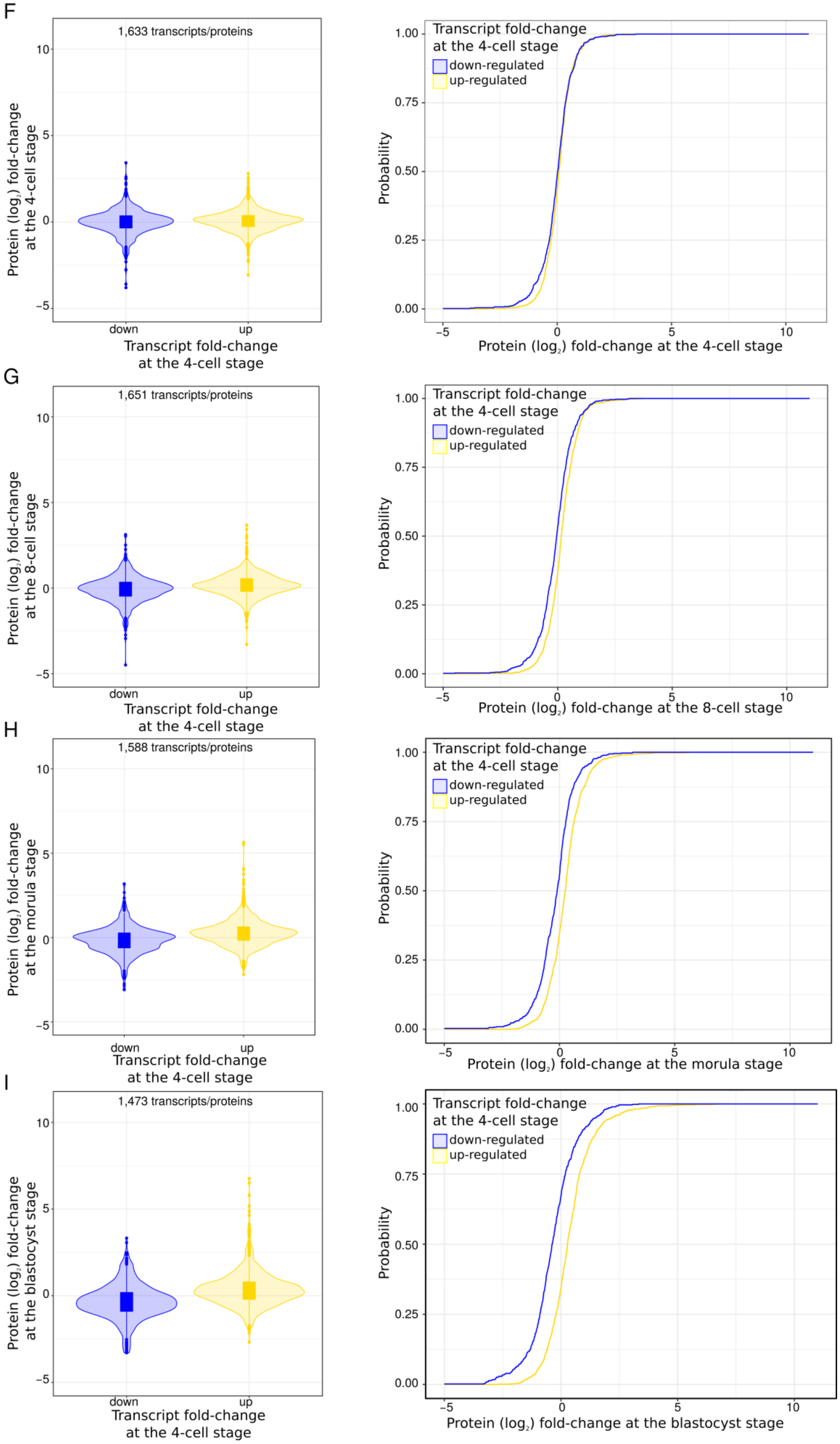
F
1,633 transcripts/proteins
Protein (log2) fold-change at the 4-cell stage
Transcript fold-change at the 4-cell stage
down
up
Transcript fold-change at the 4-cell stage
down-regulated
up-regulated
Probability
Protein (log2) fold-change at the 4-cell stage
G
1,651 transcripts/proteins
Protein (log2) fold-change at the 8-cell stage
Transcript fold-change at the 4-cell stage
Protein (log2) fold-change at the 8-cell stage
H
1,588 transcripts/proteins
Protein (log2) fold-change at the morula stage
Transcript fold-change at the 4-cell stage
Protein (log2) fold-change at the morula stage
I
1,473 transcripts/proteins
Protein (log2) fold-change at the blastocyst stage
Transcript fold-change at the 4-cell stage
Protein (log2) fold-change at the blastocyst stage

J

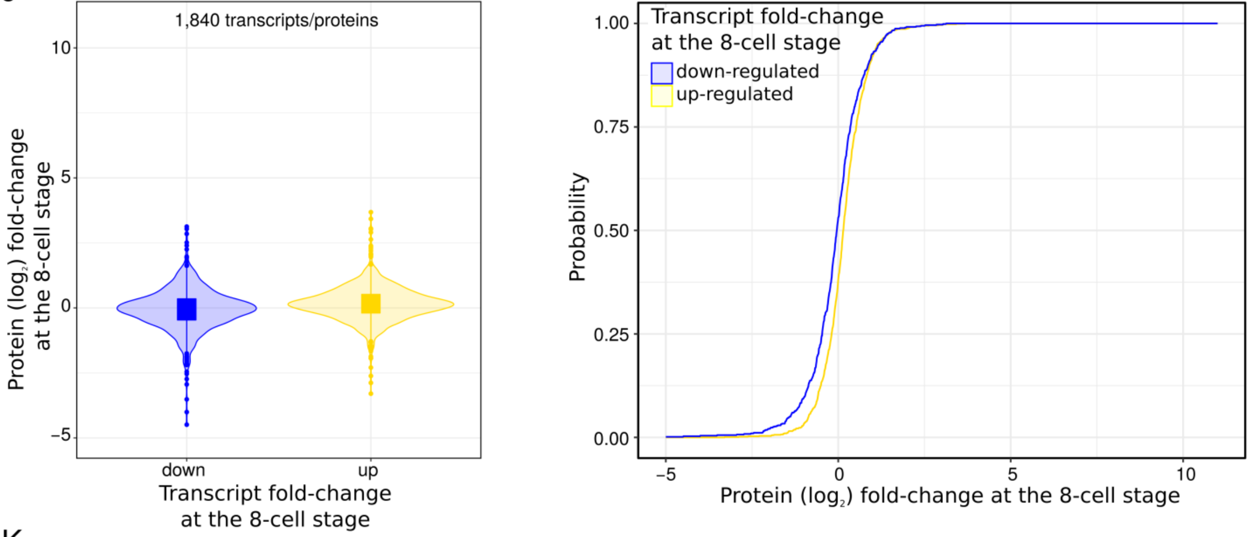

1,840 transcripts/proteins
Protein (log₂) fold-change at the 8-cell stage
Transcript fold-change at the 8-cell stage
down
up
10
5
0
−5
Transcript fold-change at the 8-cell stage
down-regulated
up-regulated
Probability
1.00
0.75
0.50
0.25
0.00
−5
0
5
10
Protein (log₂) fold-change at the 8-cell stage

K

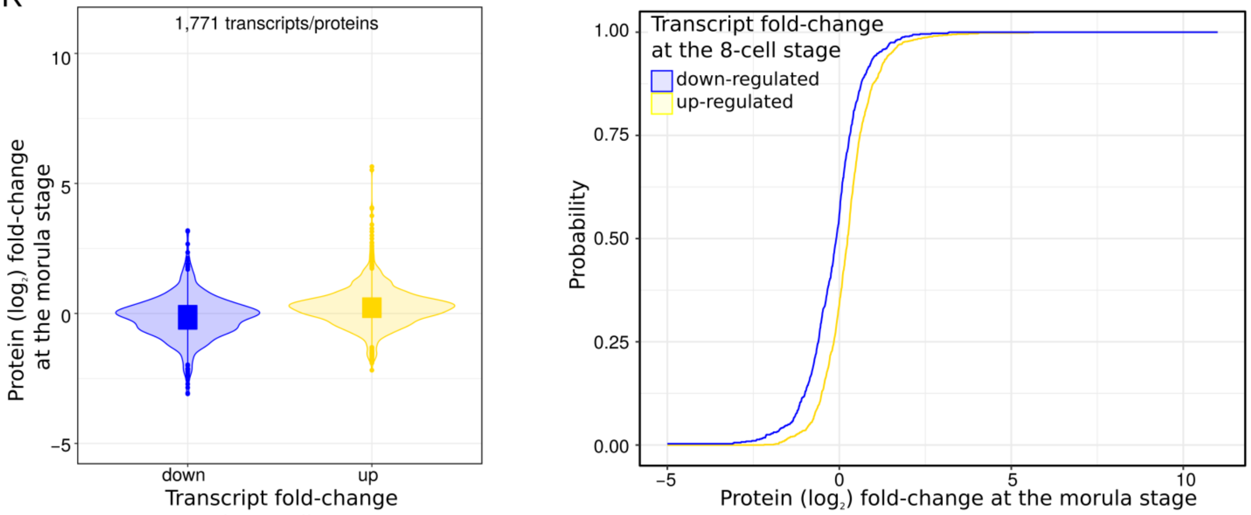

1,771 transcripts/proteins
Protein (log₂) fold-change at the morula stage
Transcript fold-change at the 8-cell stage
down
up
10
5
0
−5
Transcript fold-change at the 8-cell stage
down-regulated
up-regulated
Probability
1.00
0.75
0.50
0.25
0.00
−5
0
5
10
Protein (log₂) fold-change at the morula stage

L

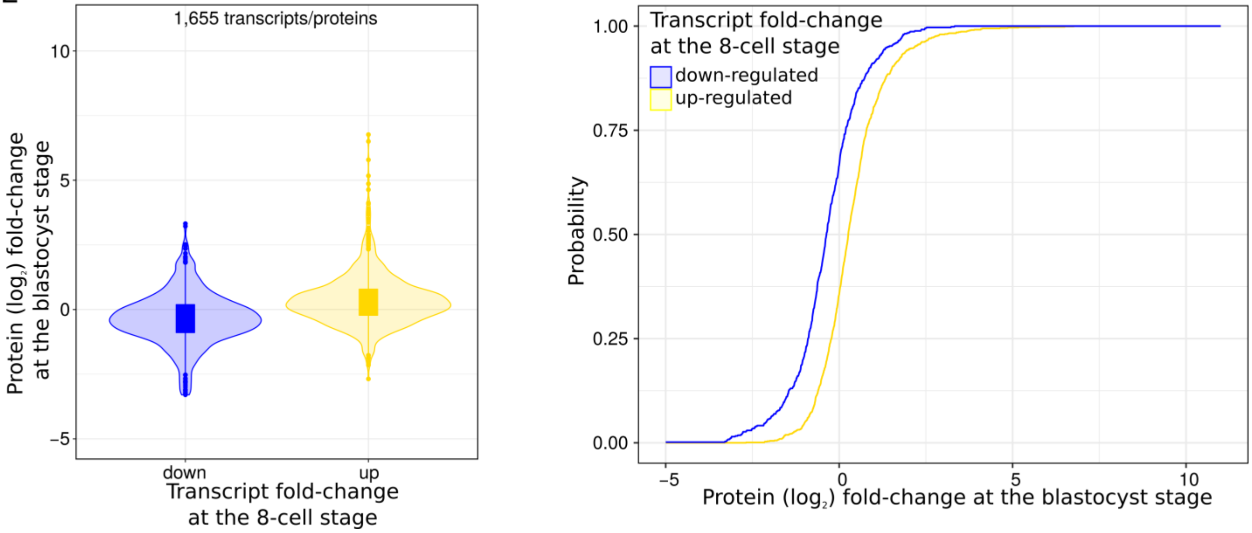

1,655 transcripts/proteins
Protein (log₂) fold-change at the blastocyst stage
Transcript fold-change at the 8-cell stage
down
up
10
5
0
−5
Transcript fold-change at the 8-cell stage
down-regulated
up-regulated
Probability
1.00
0.75
0.50
0.25
0.00
−5
0
5
10
Protein (log₂) fold-change at the blastocyst stage

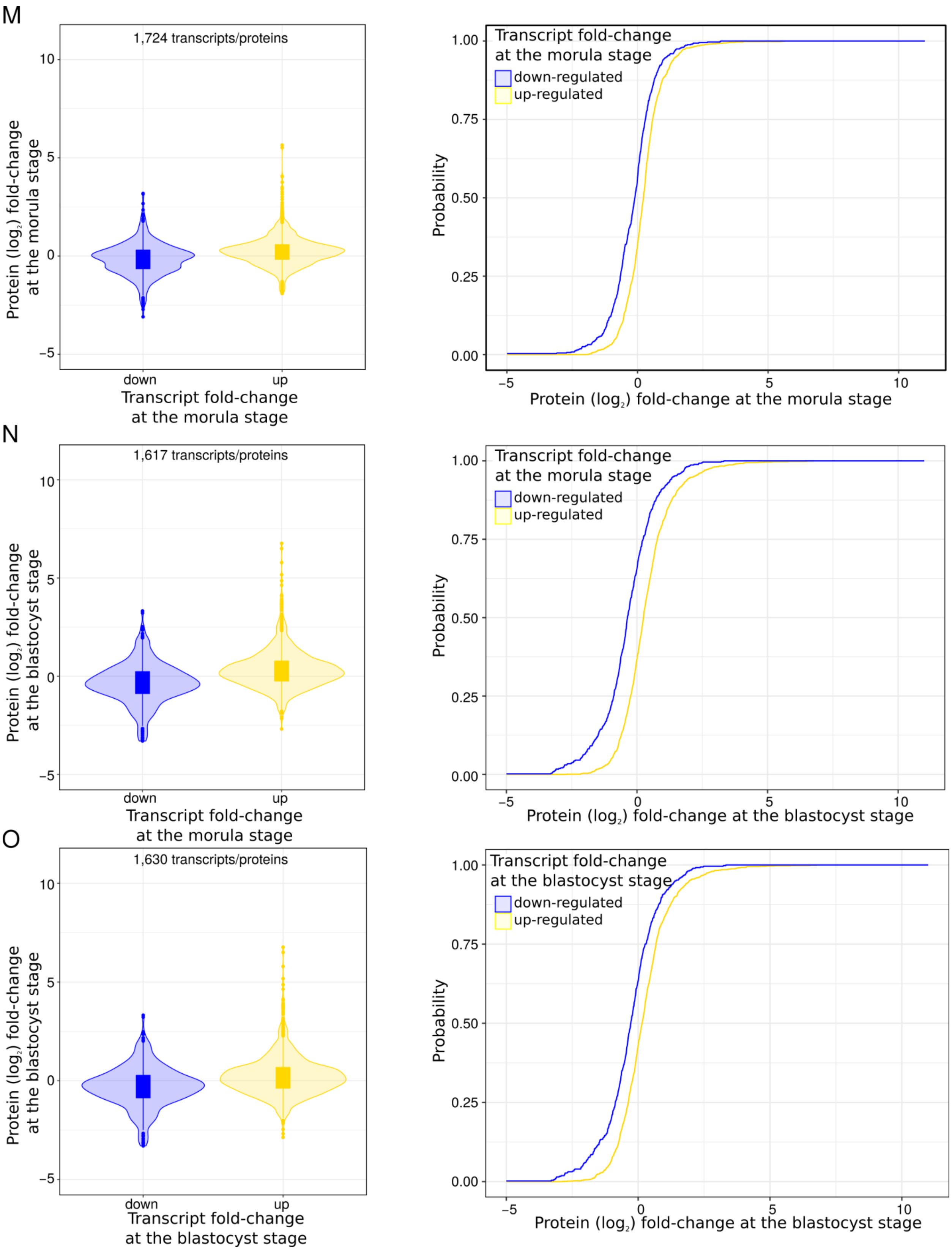

**Supplemental Fig. S16. Changes in transcriptome are only globally reflected at the proteome level from the 16-cell stage onwards.** Each panel (A-O) shows for a pair of developmental stages ($S_i$, $S_j$) the distribution of protein ($\log_2$) fold-changes at stage $S_j$ relative to the oocyte, for proteins whose cognate transcripts are down- (blue) or up- (yellow) regulated at stage $S_i$, relative to the oocyte. Left panels: Violin plots showing the distribution of protein ($\log_2$) fold-changes for proteins whose cognate transcripts are down- (blue) or up- (yellow). Right panels: Cumulative density functions (CDF) of the estimated density functions shown in left figure. Note, that in panels D, H, K and M as well as in panels E, I, L, N and O, which correspond to $S_j$ morula and blastocyst, respectively, the CDF for the proteins whose transcripts are up-regulated is shifted to the right compared to the CDF for the proteins whose transcripts are down-regulated.

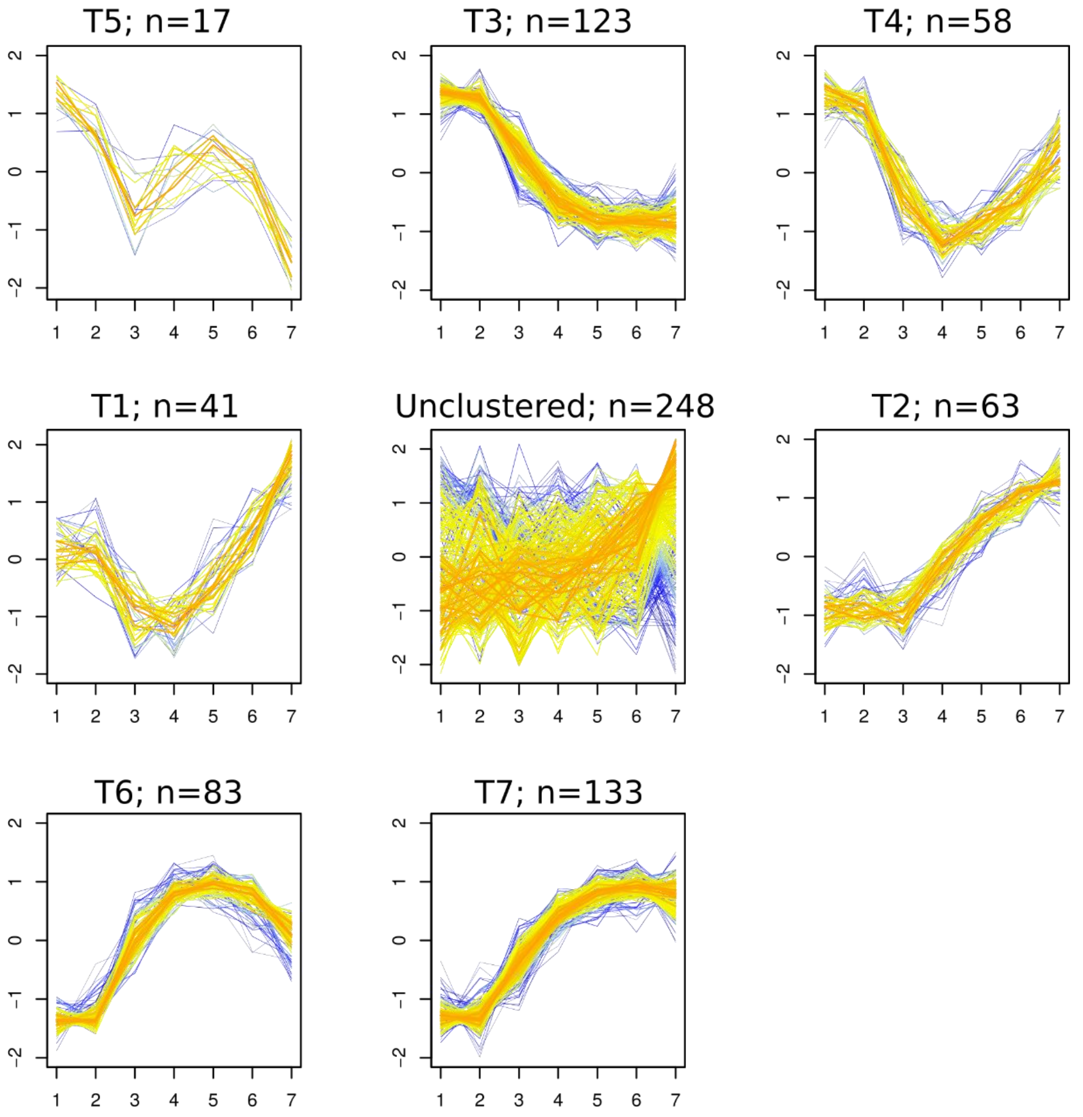

**Supplemental Fig. S17. Transcript clusters.** The x axis represents the seven developmental stages considered (1: oocyte, 2: 1-cell, 3: 2-cell, 4: 4-cell, 5: 8-cell, 6: morula and 7: blastocyst). The y axis represents the fold-change relative to the oocyte, expressed in terms of standard deviation. The coloring reflects the similarity of the temporal profile of a protein to the median profile of its cluster (red: more similar; blue: less similar).

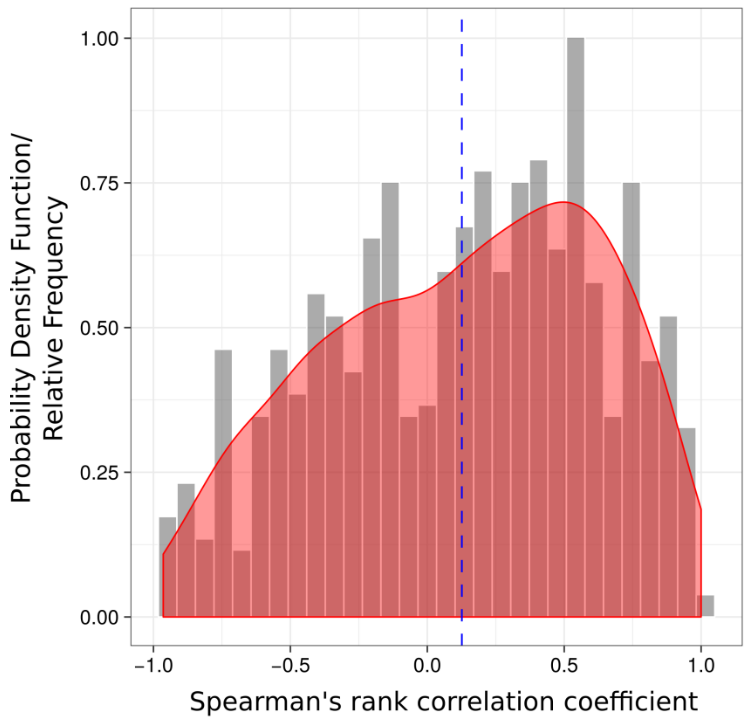

**Supplemental Fig. S18. Scaled relative frequency histogram (gray bars) and estimated probability density function (red) for the Spearman's rank correlation coefficients computed between the transcript regularized log-transformed read counts and protein L/H ratios across seven developmental stages for each gene.** Seven hundred seventy-two genes corresponding to differentially expressed proteins were considered for this analysis. The blue dashed line shows the median.

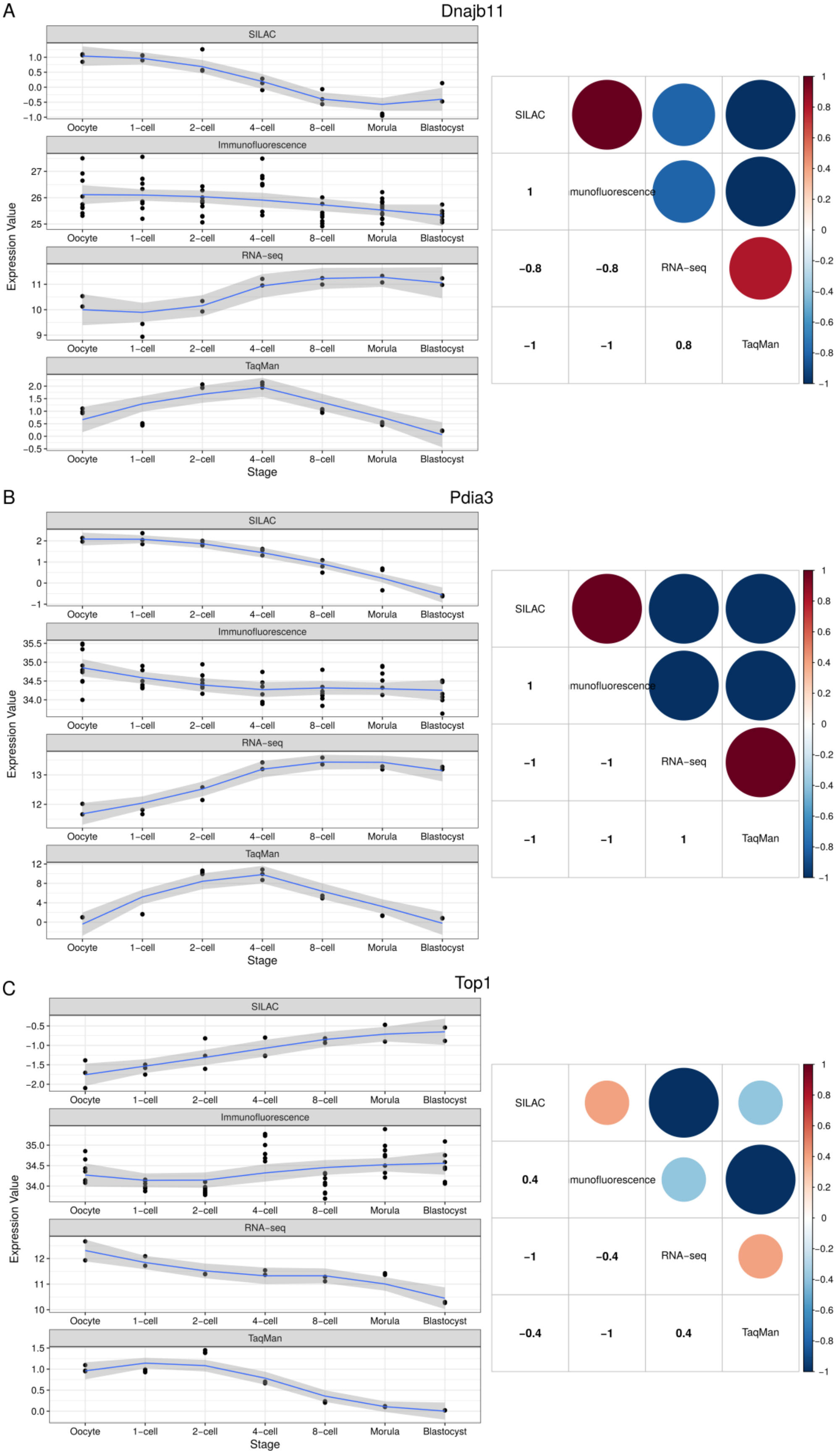
A
Dnajb11
SILAC
Immunofluorescence
RNA-seq
TaqMan
Expression Value
Oocyte
1-cell
2-cell
4-cell
8-cell
Morula
Blastocyst
Stage
SILAC
1
munofluorescence
-0.8
-0.8
RNA-seq
-1
-1
0.8
TaqMan
B
Pdia3
SILAC
Immunofluorescence
RNA-seq
TaqMan
Expression Value
Stage
SILAC
1
munofluorescence
-1
-1
RNA-seq
-1
-1
1
TaqMan
C
Top1
SILAC
Immunofluorescence
RNA-seq
TaqMan
Expression Value
Stage
SILAC
0.4
munofluorescence
-1
-0.4
RNA-seq
-0.4
-1
0.4
TaqMan

**Supplemental Fig. S19. Validation of the SILAC and RNA-seq data using immunofluorescence and TaqMan assays, respectively.** (left) Expression in the oocyte and six developmental stages. Data variation was smoothed using loess (with a span parameter of 1) curves (in blue) across the ranks corresponding to the developmental progression of the embryo starting from the oocyte, with 95% confidence interval (in grey). (right) Spearman's rank correlation coefficients (lower half) computed between the fitted loess protein/transcript profiles across the oocyte, 1-, 2- and 4-cell embryos as determined by SILAC, immunofluorescence, RNA-seq, and TaqMan assays. Transcriptomic measurements for early and late 2-cell stages were averaged. The correlation matrix was visualized using the R corrplot package (Wei and Simko 2017). The size and color of the circles (upper half) are both indicators of the magnitude of the correlation.

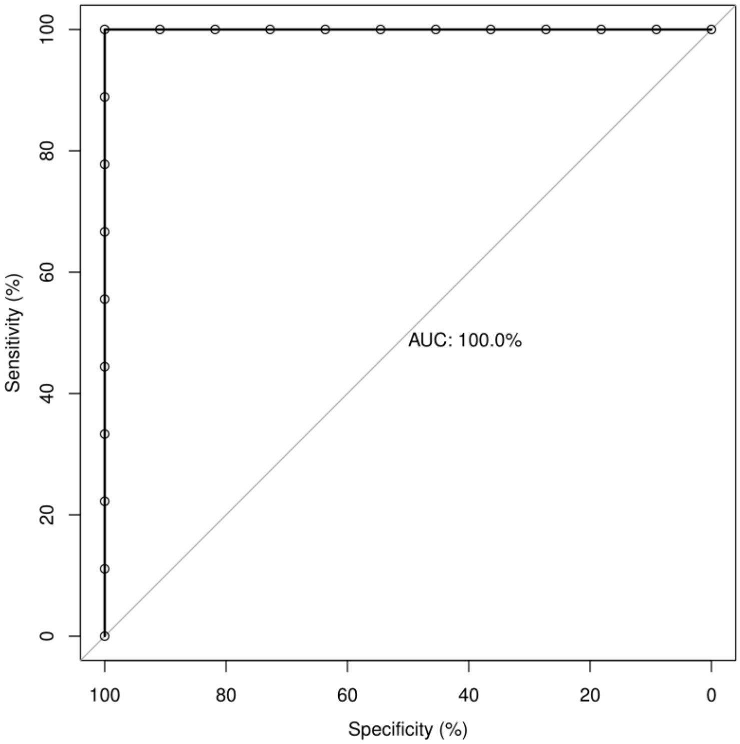

**Supplemental Fig. S20. Prediction (binary classification) of early versus late developmental stages based solely on the protein expression values. Receiver Operating Characteristic (ROC) curves (area under the curve [AUC] value is indicated).** The performance of the classifier was evaluated in a leave-one-out cross-validation (LOOCV) framework to reduce the risk of overfitting. The ROC was constructed using the posterior probabilities of the LDA for each of the test samples. Each point on the ROC curve represents a sensitivity/specificity pair corresponding to the posterior probability of a particular test sample. The AUC of 1.00 indicates that the LDA classifier is perfectly able to separate early from late developmental stages.

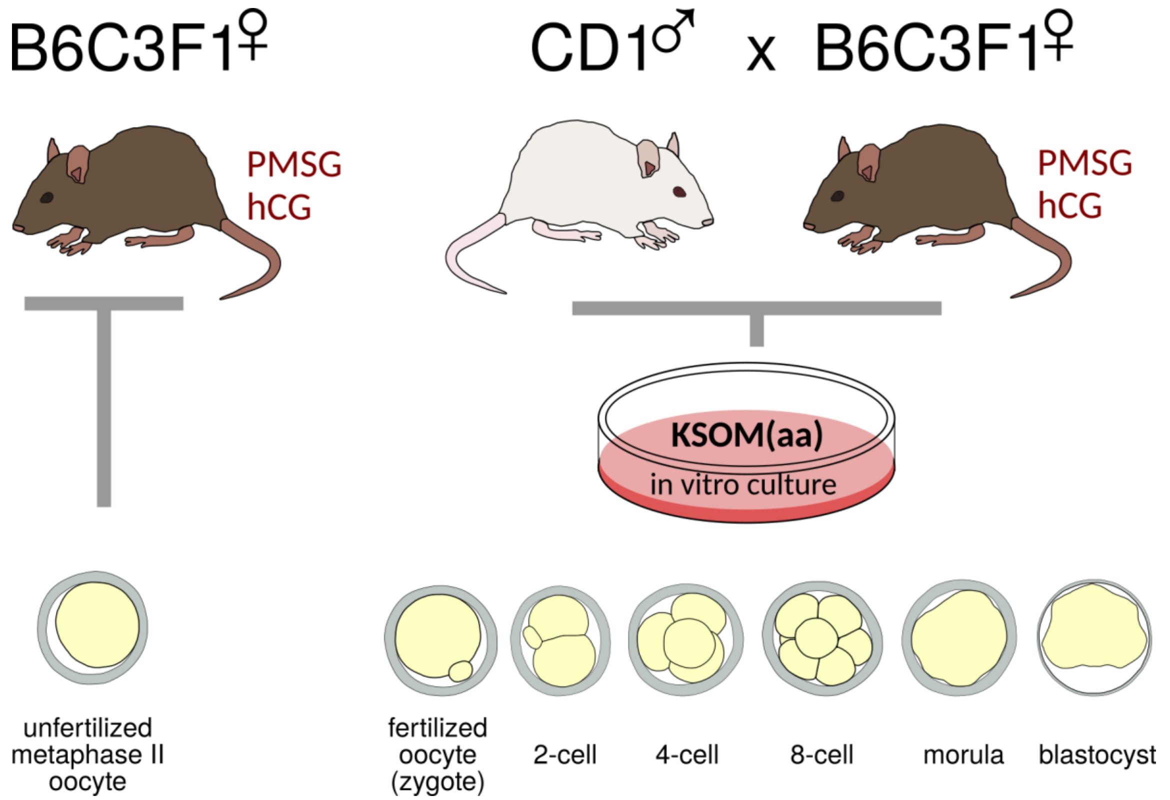

**Supplemental Fig. S21. *In vivo*-fertilized, *in vitro*-cultured mouse oocytes as a source of embryonic material for proteomic analysis.** Metaphase II or pronuclear-stage oocytes were collected from the oviducts of superovulated females, and the latter were cultured in KSOM (Potassium simplex optimized medium) medium with aminoacids to collect embryonic stages at specific time points, up to blastocyst. PMSG, pregnant mare's serum gonadotropin. hCG, human chorionic gonadotropin.

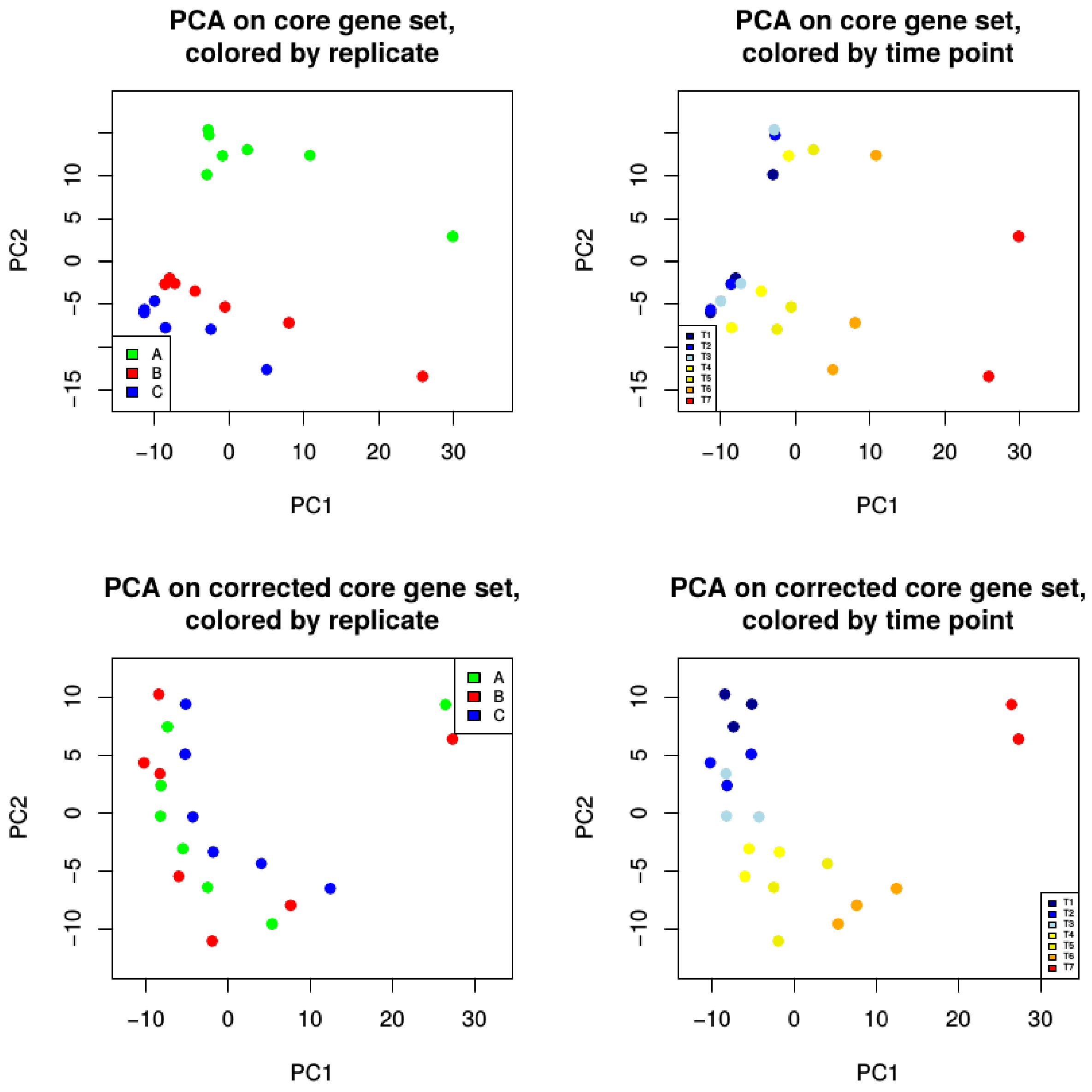

**Supplemental Fig. S22. Protein dataset batch correction. Principal Component Analysis (PCA) on the proteome dataset, based on proteins detected in all replicates of all developmental stages.** The upper panels visualize the first two PCs before applying batch correction, the lower panels visualize them after applying batch correction. In the left panels, samples are colored according to the replicate they belong to, in the right panel they are colored according to stage (T1: oocyte; T2: 1-cell; T3: 2-cell; T4: 4-cell; T5: 8-cell; T6: morula; T7: blastocyst). In the upper panels, the high contribution of the replicate identity on total variation is clearly visible, motivating the application of a batch correction procedure, resulting in a corrected visualization of the PCs that reflects the temporal progression from the oocyte to the blastocyst stage.

### Supplemental Tables

**Supplemental Table S1. Functional analysis of proteins that are differentially expressed between pairs of consecutive developmental stages ($\log_2$ fold-change ≥1 or ≤−1 between any two developmental stages, P-value≤0.05 from ANOVA).** The detected proteome was used as background for the analysis. Significantly enriched (FDR-adjusted P-value<5%) gene ontology terms (GOTERM), pathways (KEGG_PATHWAY) and keywords (KEYWORDS) were identified with DAVID (http://david.abcc.ncifcrf.gov/). The columns in the table contain information from the DAVID Functional Annotation Chart Report: Category (source provenance for the Term); Term (gene set name); Count (number of genes associated with this gene set); P-value (modified Fisher Exact P-value); List Total (number of genes in your query list mapped to any gene set in this ontology); Pop Hits (number of genes annotated to this gene set on the background list); Pop Total (number of genes on the background list mapped to any gene set in this ontology); and FDR.

**Supplemental Table S2. Functional analysis of the protein clusters.** The detected proteome (see Supplemental Table S5) was used as background for the analysis.

**Supplemental Table S3. Immunofluorescence and enzymatic validation of protein abundances.**

**Supplemental Table S4. Functional analysis of the transcript clusters.** The detected transcriptome was used as background for the analysis.

**Supplemental Table S5. Batch-corrected quantile-normalized protein L/H ratios for all samples in this study.**
