## Supplementary material for "An integrated genome-wide multi-omics analysis of gene expression dynamics in the preimplantation mouse embryo"

**Running Title:** Multi-omics of the preimplantation mouse embryo

**Keywords:** Preimplantation development, Proteome, Transcriptome, Model Organism

Supplementary Methods .......... 3
References .......... 12

### Supplementary Methods

#### MS/MS spectra search against mouse protein database

MS/MS spectra were searched against the mouse UniprotKB database (version from Dec. 2015, (The UniProt 2017)) concatenated with reversed sequence versions of all entries and supplemented with common contaminants. Parameters defined for the search were trypsin as the digesting enzyme, allowing two missed cleavages; a minimum length of seven amino acids; carbamidomethylation at cysteine residues as fixed modification, oxidation at methionine and protein N-terminal acetylation as variable modifications. The maximum allowed mass deviation was 20 ppm for the MS and 0.5 Da for the MS/MS scans. Protein groups were identified with a false discovery rate set to 1% for all peptide and protein identifications separately, when there were at least two matching peptides, at least one of which was unique to the protein group.

#### Proteome and transcriptome clustering

First, for each protein detected at least at two developmental stages in at least two replicates we computed a linear model:

$$\log_2 \frac{L}{H} = \mu + T_i + \epsilon$$

where $\mu$ is the global mean for the gene, $T_i$ is a categorical explanatory variable representing the developmental stage, and $\epsilon$ denotes the error. Next, we used the ANOVA P-value corresponding to $T_i$ as filtering criterion, and retained only those proteins for which P-value $\leq$ 0.05. Additionally, we required the proteins to have a difference of at least 2 between the largest and the smallest $\log_2 \frac{L}{H}$ ratio across all developmental stages. This resulted in 764 proteins.

We clustered the ($\log_2$) fold-change of the protein L/H ratios relative to the oocyte. ($\log_2$) fold-changes were centered and scaled before performing clustering with the R package Mfuzz (Futschik and Carlisle 2005; Kumar and M 2007). For each resulting cluster we then constructed a graph with nodes representing the members of the cluster and edges representing Pearson correlation coefficients between the expression profiles of the proteins associated with the nodes such that $r^2 \geq 0.5$. Finally, only those proteins within complete subgraphs (cliques) were selected as final members of the cluster.

We clustered the cognate transcripts of the aforementioned 764 proteins into seven clusters following an analogous procedure. The 2-cell stage represents the average of the ($\log_2$) fold-changes between the early and late 2-cell stages.

Overlap between protein and transcript clusters

The significance of the overlap between the members of all possible pairs of protein and transcript clusters (based on their official gene symbols) was computed using a Fisher's exact test with the 764 proteins/transcripts initially subjected to clustering serving as background.

**Functional enrichment analysis**

Gene Ontology (GO) enrichment analysis was performed using the Database for Annotation and Integrated Discovery (DAVID, Version 6.8, (Huang da et al. 2009b; Huang da et al. 2009a)). Proteins and transcripts were submitted to DAVID using Ensembl IDs specifying *Mus musculus* as the species. Significantly overrepresented biological process (BP), molecular function (MF) and cellular component (CC) GO terms were retrieved by using the options GOTERM_BP_ALL, GOTERM_BP_DIRECT, GOTERM_BP_FAT, GOTERM_CC_ALL, GOTERM_CC_DIRECT, GOTERM_CC_FAT, GOTERM_MF_ALL, GOTERM_MF_DIRECT, GOTERM_MF_FAT, KEGG_PATHWAY, UP_KEYWORDS, and UP_TISSUE. The default parameters and corresponding false discovery rate (FDR) by the Benjamini and Hochberg approach (Benjamini and Hochberg 1995) were used to determine significant enrichment.

In this context, we refer to the theoretical proteome/transcriptome as the entire potential protein complement encoded by the genome of the mouse, and distinguish it from the detected proteome/transcriptome, comprising only the proteins/transcripts detected in at least one replicate. For each pair of protein and transcript clusters, GO term associations for shared proteins/transcripts were tested for significance against a background set consisting of the members of the corresponding protein cluster.

**Concordance between the proteome and the transcriptome**

For every pair of developmental stages $S_i$ and $S_j$ we separated the proteins into two groups according to the change in expression (up- or down-regulation) of their cognate transcripts at $S_i$ relative to the oocyte. At each developmental stage, only proteins detected in at least two of the replicates and of the oocyte and their corresponding transcripts were considered. A gene was considered up-regulated if it was significantly differentially expressed (see Methods) and exhibited a fold-change $\geq 2$; on the contrary, a gene was considered down-regulated if it was significantly differentially expressed (see Methods) and exhibited a fold-change $\leq 0.5$. For each of the two resulting groups of proteins, we estimated the cumulative distribution function (CDF) of their ($\log_2$) fold-changes at $S_j$ relative to the oocyte using the ecdf() function in R. Thus, for each pair of developmental time points, we estimated two such CDFs: one for the proteins whose transcripts are up-regulated and one for the proteins whose transcripts are down-regulated. Finally, we integrated the two CDFs between the minimum and maximum ($\log_2$) fold-changes as determined above, yielding two areas under the curve (AUCs). To quantify the shift between the CDFs we subtracted the AUC for the proteins whose transcripts are up-regulated from the AUC for the proteins whose transcripts are down-regulated. For robustness, only sets containing at least 25 proteins/transcripts were used for calculating the AUC. This approach is common in other contexts, such as drug discovery, and has been used to describe the relationship between the transcriptome and the proteome, for example, in the ageing rat (Ori et al. 2015). For the transcriptome, the 2-cell stage represents the average of the ($\log_2$) fold-changes between the early and late 2-cell stages; a transcript is considered differentially expressed at the 2-cell stage if it is differentially expressed at the early or late 2-cell stage relative to the oocyte.

#### Number of clusters

Note that the number of clusters $k$ is a parameter of the algorithm. We therefore performed fuzzy clustering for $k \in \{4, 5, 6, 7, 8\}$ and examined the resulting membership matrices, which specify the probability with which each protein/transcript is assigned to each cluster. Specifically, we computed a PCA on this matrix using the clusters as features. We then determined the members of each cluster (based on the hard-clustering) and computed an ellipse (using the function ellipse() in the R package ellipse with default parameters) defined by the covariance matrix of the cluster PC scores and centered on the mean of the PC scores. Finally, we visually inspected the PCA plot derived from the membership matrix with the ellipses for each cluster overlaid on top. We chose the largest value of $k$ for which the ellipses are clearly separated from each other (see Figures below).

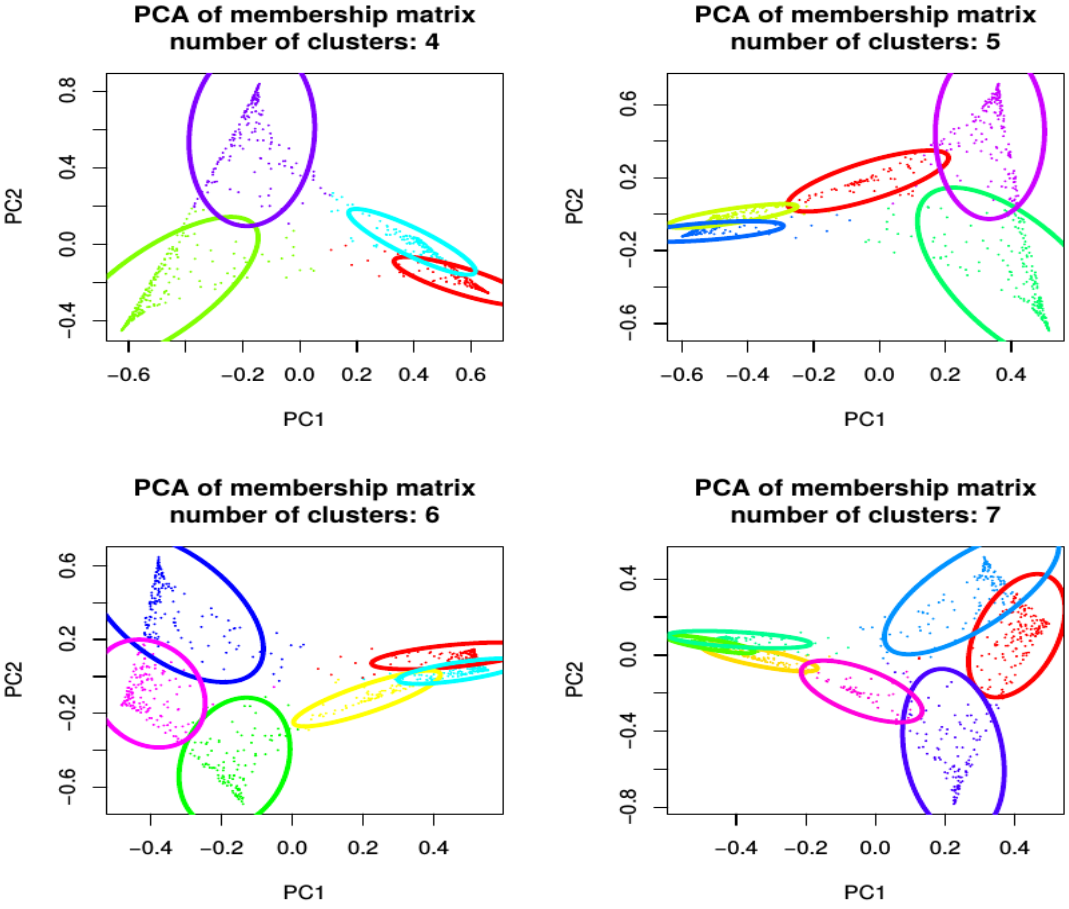

**Number of protein clusters.** We performed fuzzy clustering on the protein profiles. Like k-means, the fuzzy clustering algorithm requires the number of clusters *k* to be set in advance. We performed clustering for *k*=4, 5, 6 and 7 and principal component analysis (PCA) on each of the resulting membership matrices. The figures show the first two PCs. The ellipses are based on the mean and the covariance matrix of each cluster. A clear separation between the ellipses is indicative of distinct clusters. We chose the highest *k* for which the ellipses were clearly separated, in this case, 6. Note than when enforcing seven clusters (bottom right panel), the seventh one is wedged in between two preexisting ones (upper left portion of the subfigure, note that orientation and coloring are arbitrary, thus not comparable between subfigures).

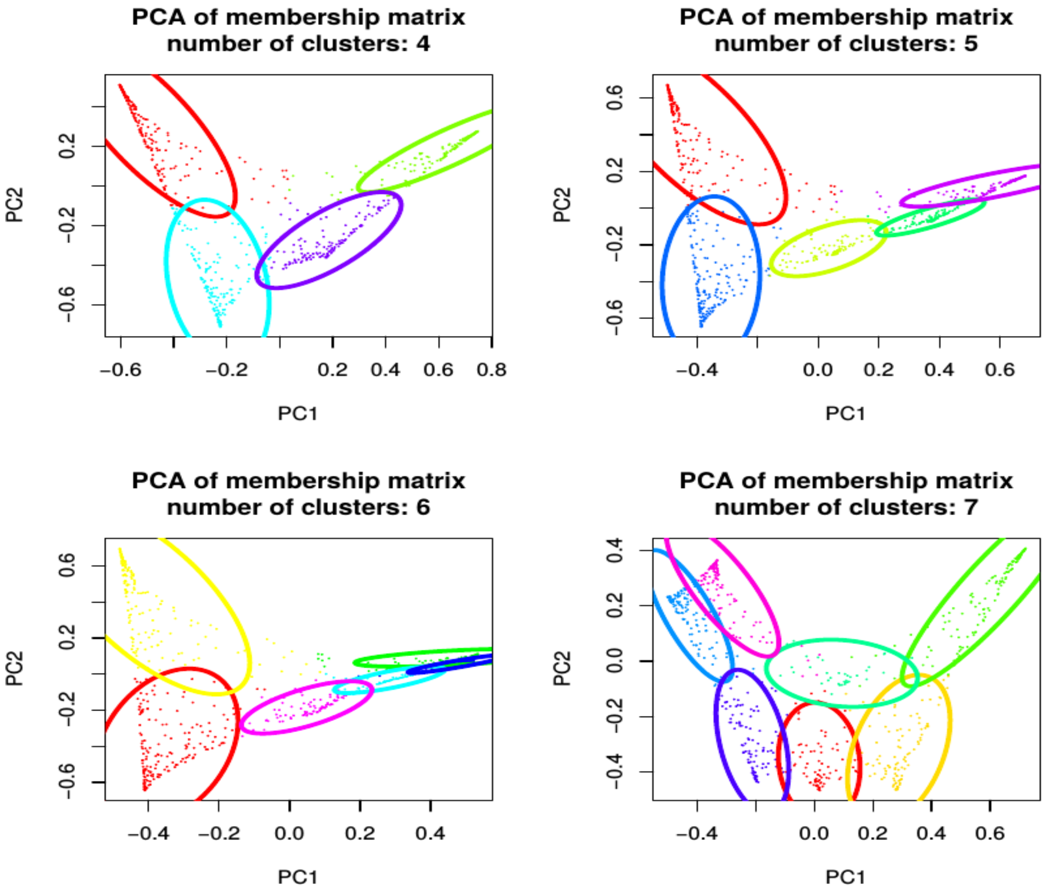


**Number of transcript clusters.** See figure above for details. Best separation is observed for 7 clusters.

#### Cluster profiles

The profile of each cluster was defined by computing the median expression value across its members for each developmental stage.

#### Markers for early and late preimplantation developmental stages

This analysis was performed using the proteins detected in all replicates of all developmental stages (1,709 proteins). Given $N_1$ samples from sample group $S_i$ and $N_j$ samples from sample group $S_j$, the classification problem is to predict the sample group (class) of any other sample based only on its protein $\log_2$ L/H ratios. For the evaluation of the classification models we used leave-one-out cross-validation, meaning that at each iteration of the cross-validation procedure, each of the $N_i + N_j$ samples was used for testing exactly once, while the remaining $N_i + N_j - 1$ samples were used for training. Thus, for a pair of sample groups $S_i$ and $S_j$, we trained and tested $N_i + N_j$ models. For producing the results reported here we grouped the developmental stages into two sample groups: (i) early (oocyte, 1- and 2-cell-embryo stages, $N_1 = 3 + 3 + 3 = 9$) and late preimplantation development (4- and 8-cell, morula and blastocyst embryo stages, $N_1 = 3 + 3 + 3 + 2 = 11$), and trained a total of 20 classifiers. The reported classification rate is the fraction of correctly classified samples among the 20 tests.

At each iteration of the cross-validation procedure, we selected features with a mean $\log_2$ L/H ratio $\geq 1$ in at least one of the two classes in the training set and discarded the rest from further analysis. Next, we performed Principal Component Analysis (PCA). To regularize the classification problem and avoid over-fitting of the classifier, only the first two principal components (PC1 and PC2) were used as features to train a Linear Discriminant Analysis (LDA) model. The aim of LDA is to find a hyperplane that separates between two classes based on their features.

To classify the test sample we first projected its $\log_2$ L/H ratios onto PC1 and PC2 by multiplying the zero-centered $\log_2$ L/H ratios with the PCA rotation matrix and keeping only the first two components, and assigned a class based on the aforementioned hyperplane. If the LDA did not achieve a perfect class separation for the training sample, the classification of the test sample was considered “incorrect” and the classification rate computed accordingly.

Finally, we identified the features that were among the most discriminant features at all iterations of the cross-validation procedure. A protein ranking was thus extracted based on the LDA. Specifically, we projected PC1 and PC2 onto the LDA space. This projection is a linear combination of PC1 and PC2, which are in turn linear combinations of the original protein $\log_2$ L/H ratios. Hence, the most discriminant proteins are those with the highest and lowest coefficients in the linear combination. We computed a rank for each classifier and aggregated them using the RankAggregg() function in the RankAggregg R package (Pihur et al. 2009). From this aggregated ranking, we selected the top 20 proteins as the most discriminant features between the two classes.

#### Comparison to other published datasets

To compare our dataset to that of Gao et al. (Gao et al. 2017) we first used the org.Mm.eg.db R/BioConductor package (version 3.4.1, (Carlson 2018)) to identify (and translate, if necessary) the official symbol of the proteins included in Gao et al.'s Table S1. In approximately 10% of the cases, the reported symbols were aliases. For the sake of compatibility, we excluded our oocyte data from the comparison.

#### Subcellular localization of protein complexes

The definition of the protein complexes was obtained from the resource compiled by Ori et al (Ori et al. 2016). Specifically, we downloaded Additional file 2, which comprises the descriptions of 279 protein complexes, including the human Ensembl gene identifier of their members. Human Ensembl gene identifiers were mapped to mouse orthologs using the Ensembl orthology data (Vilella et al. 2009; Flicek et al. 2014) and the BioMart interface of Ensembl (Kinsella et al. 2011). Out of a total of 279 protein complexes, 233 had 5 or more mouse members and were considered for the analysis.

The (preferential) subcellular localization(s) of the protein complexes were determined based on the annotation of the COMPARTMENTS database (https://compartments.jensenlab.org/, (Binder et al. 2014)). Specifically, each complex was assigned the subcellular localization(s) of the largest group(s) of proteins among its members, as annotated in the Compartments database. The vast majority of the complexes (199/233) were associated with only one subcellular localization.

#### Classification into structural proteins, enzymes and transcription factors

Protein groups were defined based on the PANTHER Classification System (http://www.pantherdb.org/, (Mi et al. 2013; Mi et al. 2017)), which has a parent/child hierarchical organization of the corresponding database. The database was queried by means of the Bioconductor/R package PANTHER.db (Muller). Thus:

- Structural proteins included all proteins in the PANTHER classes: "cytoskeletal protein (PC00085)", "cell junction protein (PC00070)", "structural protein (PC00211)", "membrane traffic protein (PC00150)", "extracellular matrix protein (PC00102)", "cell adhesion molecule (PC00069)" and "viral coat protein (PC00236)" and all their children/descendants. A total of 1,875 proteins in the PANTHER database satisfied this condition.
- Enzymes are all proteins in PANTHER classes with the “-ase” suffix (e.g., protein kinase, phospholipase, oxidoreductase) and their children/descendants. A total of 3,988 proteins in the PANTHER database satisfied this condition.
- Transcription factors comprised all proteins in the PANTHER class “transcription factors” (PC00218) and all its children/descendants. A total of 1,392 proteins in the PANTHER database satisfied this condition.
